# EF-Hand Calcium Sensor, EfhP, Controls Transcriptional Regulation of Iron Uptake by Calcium in *Pseudomonas aeruginosa*

**DOI:** 10.1101/2024.01.09.574892

**Authors:** Jacob Burch-Konda, Biraj B. Kayastha, Aya Kubo, Myriam Achour, Mackenzie Hull, Reygan Braga, Lorelei Winton, Rendi R. Rogers, Jacee McCoy, Erika I. Lutter, Marianna A. Patrauchan

**Affiliations:** Department of Microbiology and Molecular Genetics, Oklahoma State University, Stillwater, OK 74078, USA

**Keywords:** Transcription, calcium signaling, promoter activity, pyoverdine, RNA seq, virulence, human pathogen, CarRS, BqsRS, Fur, CF clinical isolates

## Abstract

The human pathogen *Pseudomonas aeruginosa* poses a major risk for a range of severe infections, particularly lung infections in patients suffering from cystic fibrosis (CF). As previously reported, the virulent behavior of this pathogen is enhanced by elevated levels of Ca^2+^ that are commonly present in CF nasal and lung fluids. In addition, a Ca^2+^-binding EF-hand protein, EfhP (PA4107), was partially characterized and shown to be critical for the Ca^2+^-regulated virulence in *P. aeruginosa*. Here we describe the rapid (10 min, 60 min), and adaptive (12 h) transcriptional responses of PAO1 to elevated Ca^2+^ detected by genome-wide RNA sequencing and show that *efhP* deletion significantly hindered both rapid and adaptive Ca^2+^ regulation. The most differentially regulated genes included multiple Fe sequestering mechanisms, a large number of extracytoplasmic function sigma factors (ECFσ) and several virulence factors, such as production of pyocins. The Ca^2+^ regulation of Fe uptake was also observed in CF clinical isolates and appeared to involve the global regulator Fur. In addition, we showed that the *efhP* transcription is controlled by Ca^2+^ and Fe, and this regulation required Ca^2+^-dependent two-component regulatory system CarSR. Furthermore, the *efhP* expression is significantly increased in CF clinical isolates and upon pathogen internalization into epithelial cells. Overall, the results established for the first time that Ca^2+^ controls Fe sequestering mechanisms in *P. aeruginosa* and that EfhP plays a key role in the regulatory interconnectedness between Ca^2+^ and Fe signaling pathways, the two distinct and important signaling pathways that guide the pathogen’s adaptation to host.

**IMPORTANCE:** *Pseudomonas aeruginosa* (*Pa*) poses a major risk for severe infections, particularly in patients suffering from cystic fibrosis (CF). For the first time, kinetic RNA sequencing analysis identified *Pa* rapid and adaptive transcriptional responses to Ca^2+^ levels consistent with those present in CF respiratory fluids. The most highly upregulated processes include iron sequestering, iron starvation sigma factors, and self-lysis factors pyocins. An EF-hand Ca^2+^ sensor, EfhP, is required for at least 1/3 of the Ca^2+^ response, including all the iron uptake mechanisms and production of pyocins. Transcription of *efhP* itself is regulated by Ca^2+^, Fe, and increases during interactions with host epithelial cells, suggesting the protein’s important role in *Pa* infections. The findings establish the regulatory interconnectedness between Ca^2+^ and iron signaling pathways that shape *Pa* transcriptional responses. Therefore, understanding Pa’s transcriptional response to Ca^2+^ and associated regulatory mechanisms will serve the development of future therapeutics targeting *Pa* dangerous infections.

## INTRODUCTION

*Pseudomonas aeruginosa (Pa)* is a major human pathogen causing severe chronic infections. It is a primary pathogen infecting the airways of cystic fibrosis (CF) patients, leading to cellular damage and fatal obstructive lung disease (*1, 2*). *Pa* is also responsible for life threatening infective endocarditis and infections in urinary tract, skin, burn and surgical wounds (*3–6*). *Pa* infections have a mortality rate of 18 to 60% and present a tremendous health care burden in the United States (*7*) and worldwide. *Pa* belongs to the ESKAPE group of pathogens with increased virulence, persistence, and transmissibility (*8*) and has been recognized by both the Centers for Disease Control and Prevention (CDC) and the World Health Organization (WHO) as a serious threat that requires the urgent development of new therapeutics. The success of *Pa* as a pathogen and the failure of antibiotic therapies to eliminate *Pa* infections are thought to be due in part to its physiological plasticity, enabling the adaptability and resistance (reviewed in (*9, 10*)). Furthermore, *Pa* was shown to shape the host innate immune response to favor pathogen survival (*11*). However, the underlying molecular mechanisms as well as the host stimuli that trigger adaptive responses are not well understood.

Upon entry into the host, pathogenic bacteria must survive several lines of the host immune defense, which requires efficient recognition and responsiveness to an array of host messengers. To coordinate the responses to different host factors, *Pa* possesses a record number of transcriptional regulatory systems, that modulate and tune the gene expression of multiple pathways required for survival (*12, 13*). Among a multitude of host factors that control the expression of *Pa* virulence factors, calcium (Ca^2+^) is particularly influential as it also plays a major role in a human host by controlling life-sustaining processes, such as cell division, differentiation, apoptosis, gene transcription and immune responses (*14–16*). Disruption of Ca^2+^ homeostasis may cause or result from diseases, including bacterial infections and genetic disorders (*17–25*). One example is cystic fibrosis (CF), where the levels of Ca^2+^ are elevated in lungs and nasal fluids often due to imbalance of ion homeostasis (*26–30*). Cases of calcification of the lungs, vascular and extravascular regions have also been reported (*30–32*). Such elevated level of Ca^2+^ may act as a signaling cue and lead to modulation of the physiology and virulence of the invading bacteria, including *Pa* (*25, 33–37*). Therefore, a better understanding of Ca^2+^-dependent responses in *Pa* will inform about the host-pathogen relationship and help in the development of antimicrobial strategies.

Previously, we and others have collected multiple evidence illustrating the regulatory role of Ca^2+^ in *Pa.* These include Ca^2+^-induced production of virulence factors, pyocyanin (*25*) and extracellular proteases (*38, 39*), biofilm formation (*33, 40, 41*), swarming motility, and antibiotic resistance (*33, 41, 42*). These virulence factors enhance colonization by *Pa* and facilitate its evasion of the host immune system (*43–45*). In addition, elevated Ca^2+^ negatively regulates a potent virulence factor, type III secretion system (T3SS), known to promote bacterial persistence and dissemination from the infection site (*46*). In the effort to elucidate the molecular mechanisms of Ca^2+^ regulation, we have identified several key components of Ca^2+^ regulatory network in *Pa* that, among others, include the Ca^2+^-regulated two component regulatory system CarSR (*47*) (identical to BqsSR in PA14) (*48*)), Ca^2+^ channel, CalC (Guragain, etal., in preparation), and Ca^2+^ sensor, EfhP (*25, 49*).

We have established that EfhP meets all requirements of a Ca^2+^ sensor: it selectively binds Ca^2+^ and undergoes Ca^2+^-dependent conformational changes, which are required for Ca^2+^ regulation of several aspects of *Pa* virulence (*25, 49*). Disruption of *efhP* leads to a decrease in Ca^2+^-regulated production of pyocyanin, alginate, as well as *Pa* virulence in plant and *Galleria mellonella* infection models (*49, 50*). These data support a hypothesis that EfhP plays a key role in orchestrating the Ca^2+^ control over *Pa* gene expression. Furthermore, *efhP* is highly conserved in *Pseudomonas* genus with even greater sequence conservation among *Pa* CF clinical isolates vs non-CF clinical and environmental isolates (*49*), which provides additional evidence of its role in human infections. The predicted periplasmic localization of EfhP (*49*) may equip *Pa* with a mechanism to sense and respond to extracellular concentrations of Ca^2+^. Since *Pa* faces a rapid transition to millimolar levels of Ca^2+^ upon invading human airways, which are further heightened during CF, it is important to characterize the development of Ca^2+^ transcriptional responses over time. Therefore, we applied RNA sequencing to characterize the genome-wide kinetic changes in *Pa* transcriptome during adaptive and rapid responses to elevated Ca^2+^ and determine the role of EfhP in *Pa* transcriptional regulation by Ca^2+^.

## RESULTS

### The PAO1 kinetic transcriptional response to Ca^2+^ comprises a major share of the genome

Genome-wide RNA sequencing was conducted to provide a novel look at the rapid and adaptive transcriptional changes occurring in *Pa* in response to 5 mM Ca^2+^. The selected concentration of Ca^2+^ reflects those present in the nasal fluids and saliva of CF patients, where *Pa* has been shown to undergo significant patho-adaptation (*51, 52*). For the 10 min and 60 min “rapid” responses, cultures were grown to mid-log phase in the absence of Ca^2+^ before their subsequent Ca^2+^ exposures and RNA extraction. The 12 h condition was defined as the “adaptive” response as the cultures were grown in the presence of Ca^2+^ to mid-log phase before RNA extraction.

The RNAseq-based analysis of differential gene expression of the three kinetic experimental groups vs. the control PAO1 not exposed to Ca^2+^ revealed a dynamic response to Ca^2+^ (Fig. 1A). The greatest response is seen in the 10 min condition, with 1199 genes significantly upregulated (log2FC>1, p <0.05) and 1045 genes significantly downregulated (log2FC <-1, p <0.05). This rapid response settles partially by 60 min of Ca^2+^ exposure (520 genes upregulated, 446 downregulated). Following 12 h adaptation to Ca^2+^, the transcriptional response increases to 665 genes upregulated and 666 downregulated. This adaptive response includes 292 unique genes not observed to be differentially regulated in the 10 min and 60 min rapid response conditions, indicating that *Pa* utilizes a specific regulatory network to adapt to long-term Ca^2+^ exposure versus short-term (Fig. 1B). Overall, of the 5709 genes in the PAO1 genome (pseudomonas.com), 39%, 17%, and 23% were shown to be Ca^2+^-responsive in the 10 min, 60 min, and 12 h conditions respectively.

**Figure 1.**
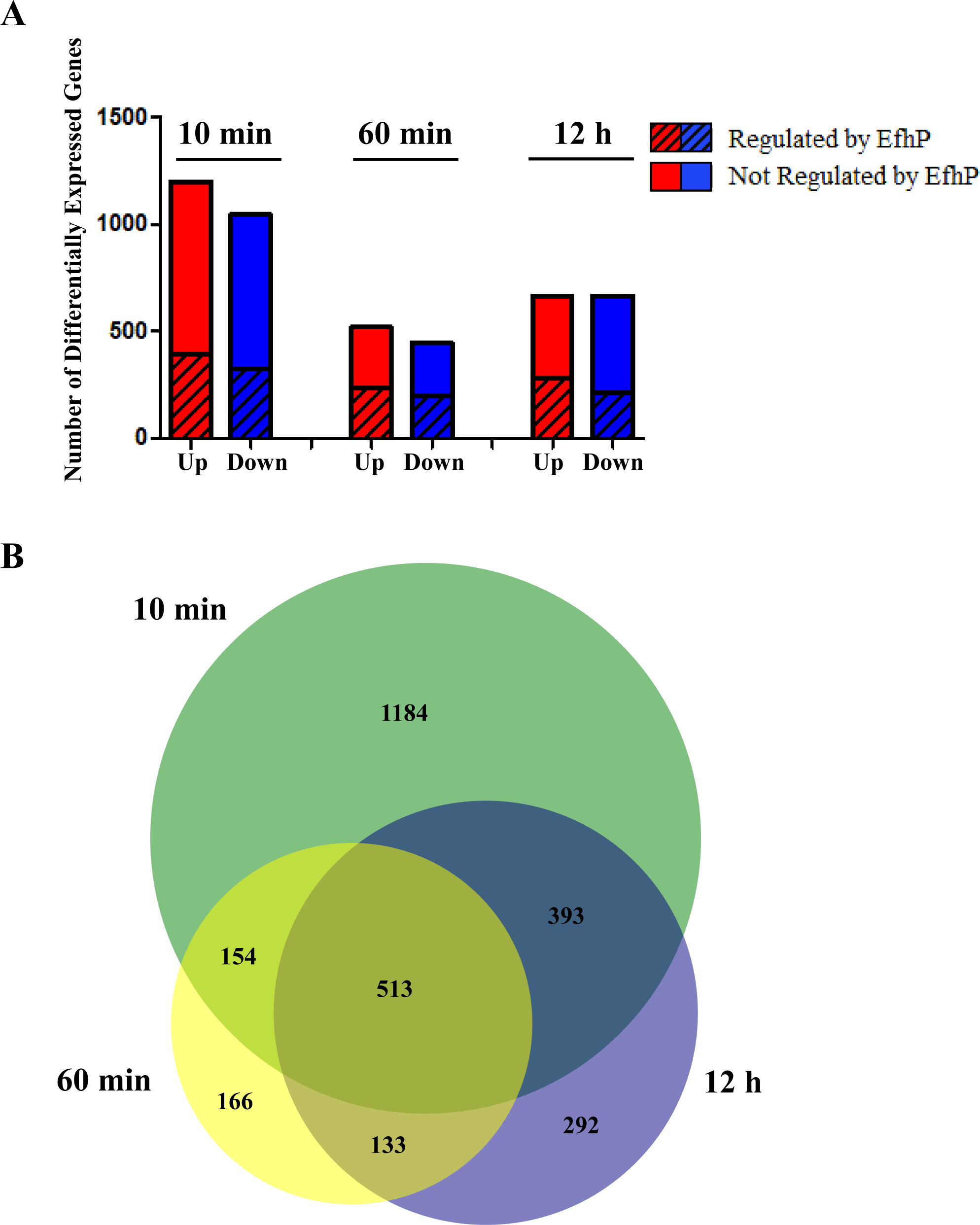
PAO1 transcriptional response to 10 min, 60 min, and 12 h exposure to Ca^2+^. **(A)** Total number of genes significantly upregulated (log2FC >1, p <0.05) and downregulated (log2FC <-1, p <0.05) at each time point are depicted in red and blue respectively. Dashes indicate the number of calcium-responsive genes that are regulated by EfhP at each time point. For PAO1 upregulated genes this refers to a log2FC< 0 in the Δ*efhP* strain in same conditions, and a log2FC> 0 for PAO1 downregulated genes. (**B)** Numbers of Ca^2+^-responsive genes (log2FC <-1 or >1, p<0.05) shared by or unique to the 10 min, 60 min and 12 h Ca^2+^ exposure conditions.

To gain a deeper understanding of the specific *Pa* functions regulated by Ca^2+^, the full PAO1 genome has been sorted into 26 functional categories (Fig. 2). These categories were assigned based on hand-curated gene annotation described in the Methods. Percent coverage of each category was calculated by dividing the numbers of differentially regulated genes by the total number of annotated genes in each category. Iron (Fe) uptake was the most Ca^2+^-responsive functional category at each time point, with 74%, 68%, and 68% coverage at the 10 min, 60 min, and 12 h time points, respectively. Following the Fe uptake genes, the next most Ca^2+^-responsive functional categories in the 10 min rapid response condition are sulfur metabolism and chemotaxis/motility. At 60 min exposure, Ca^2+^ regulation of these two categories is almost entirely suppressed, perhaps suggesting a regulatory correction by additional regulatory networks. However, Ca^2+^ retains a relatively high regulatory control of the Fe uptake, stresses (oxidative, heat, cold, osmotic) and polyamine metabolism/transport functional categories at 60 min. In the 12 h adaptive response, phosphate and phosphonate metabolism/transport and secreted factors (pyocins, phenazine, toxins) emerge as the most Ca^2+^-responsive genes after Fe uptake. The former two categories were also likely to be uniquely regulated by Ca^2+^ in the adaptive response, with 17 genes and 10 genes regulated respectively. Eight genes belonging to the Hxc type II secretion system (*53*) were additionally upregulated during the 12 h adaptive response but not at 10 or 60 min.

**Figure 2.**
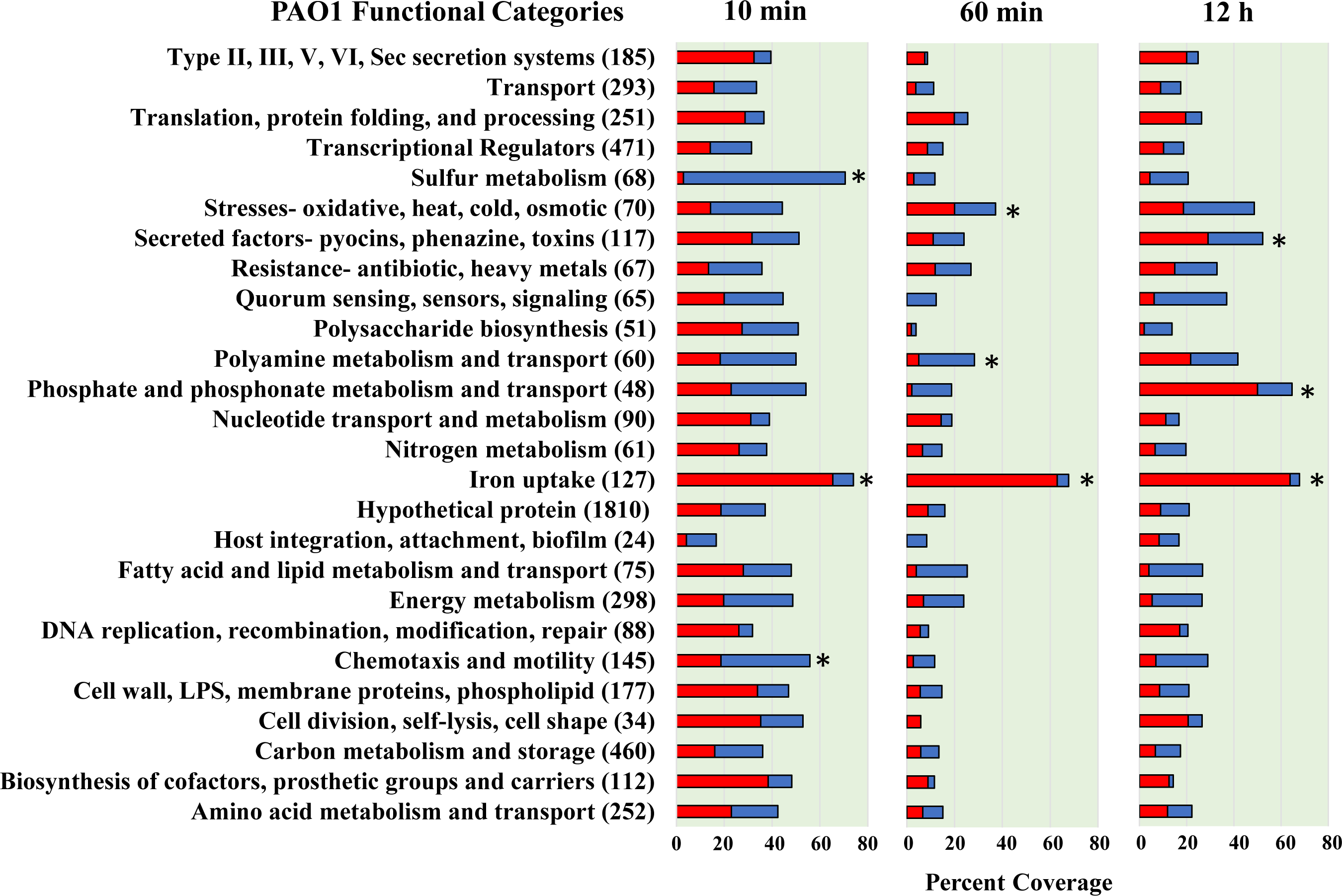
The PAO1 transcriptional response to 5mM Ca^2+^ by functional category. RNA sequencing data from samples exposed to 5mM Ca^2+^ for 10 min, 60 min, or 12 h was compared to samples grown to mid log phase in the absence of Ca^2+^. Red bars denote upregulation (log2FC >1, p <0.05) and blue bars denote downregulation (log2FC<-1, p <0.05). These data present the percent coverage of each functional category which is either significantly upregulated or downregulated at each time point. The number of genes assigned to each functional category are shown in parentheses. Functional categories were developed as described in Methods. Functional categories most highly regulated by Ca^2+^ at each time point are denoted with asterisks.

### EfhP regulates PAO1 rapid and adaptive responses to Ca^2+^

Previously, we have identified and partially characterized *Pa* Ca^2+^ sensor, EfhP (*25, 49*). Here we aimed to characterize the role of this protein in PAO1 transcriptional response to Ca^2+^. The comparative RNA-seq analysis of Δ*efhP* grown under the same conditions as PAO1 (0 min, 10 min, 60 min, and 12 h Ca^2+^ exposure) revealed that about 31% of the total Ca^2+^ response in PAO1 requires EfhP. EfhP regulation has been called when genes upregulated (log2FC >1) and downregulated (log2FC <-1) by Ca^2+^ in PAO1 showed Ca^2+^-dependent transcript changes of log2FC< 0 or log2FC> 0, respectively, in the mutant. This includes both rapid and adaptive responses (Fig. 1A). The data suggest a specific EfhP regulon within the broader PAO1 Ca^2+^ response.

### EfhP is required for Ca^2+^ regulation of iron uptake

A closer examination of EfhP’s regulation within specific PAO1 functional categories revealed that the protein is required for Ca^2+^ regulation of many pathways involved in *Pa* virulence (Fig. 3). The most striking is the EfhP-dependent regulation of Fe uptake genes in both rapid and adaptive responses to Ca^2+^ (Fig. 4A). In PAO1, over 60% of the iron uptake genes are upregulated by Ca^2+^ at all three time points, whereas in Δ*efhP* these genes either do not respond or are downregulated in response to Ca^2+^. It is important to note that samples used for RNA sequencing in this study were cultured at 3.6 µM Fe, the concentration consistent with Fe levels in the CF sputum (*54*). So, the apparent Fe starvation response developed in the presence of Fe.

**Figure 3.**
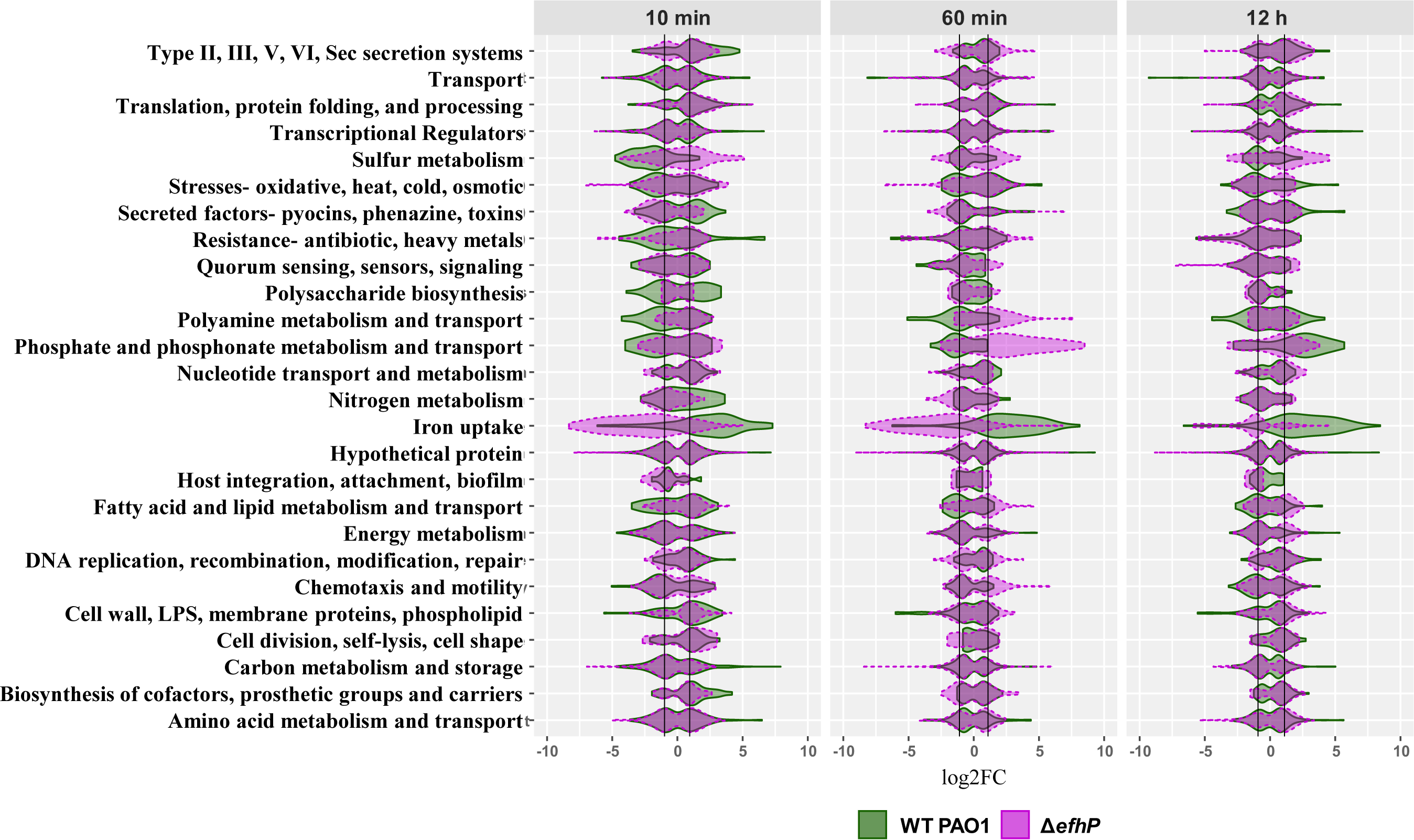
Comparing the PAO1 and Δ*efhP* transcriptional responses to 5mM Ca^2+^ by functional category. The distributions of log2FC values for genes within each functional category are plotted for both PAO1 (green) and Δ*efhP* (purple). 10 min v 0 min, 60 min v 0 min, and 12 h v 0 min condition comparisons are shown side by side. Black lines indicate log2FC= ±1. Genes falling outside of the black lines are at least twofold upregulated or downregulated. Only statistically significant genes (p < 0.05) were considered for generation of the violin plots.

**Figure 4A.**
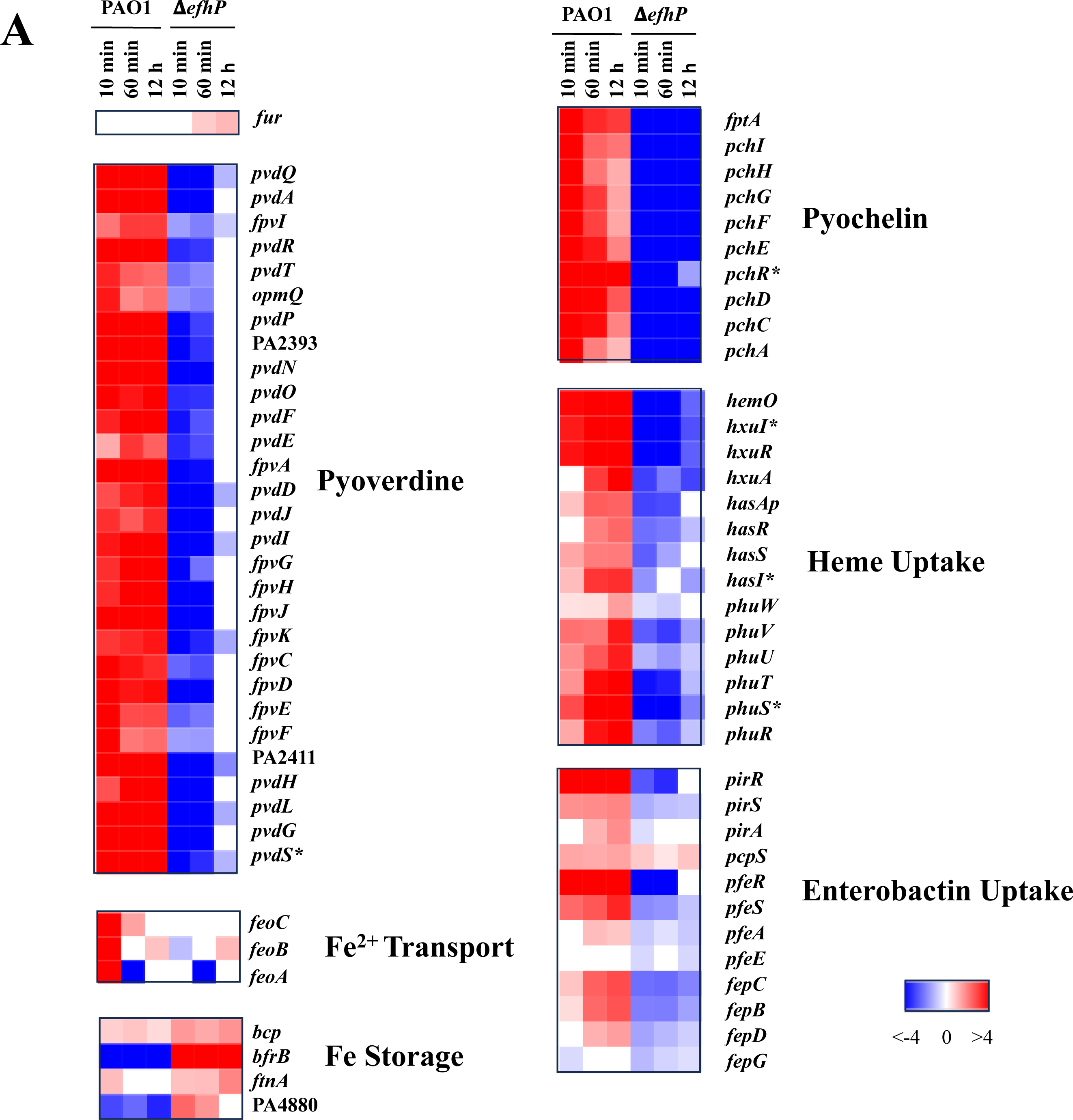
EfhP regulation of Fe uptake genes. The heat maps show log2FC values of expression comparisons of 10 min, 60 min, or 12 h 5 mM Ca^2+^ conditions vs. 0 mM Ca^2+^ condition. PAO1 and Δ*efhP* strains are compared to their respective control transcriptional profiles of cultures grown to mid log phase at 0 mM Ca^2+^. Major regulators are denoted with asterisks.

At the pathway level, the positive regulation by Ca^2+^ in PAO1 includes the genes encoding for pyoverdine, pyochelin, heme, and enterobactin uptake, along with pyoverdine and pyochelin biosynthesis genes. A rapid upregulation of these pathways was observed after 10 min and 60 min Ca^2+^ exposure in PAO1 and extends to the 12 h exposure. In contrast, most of these genes showed downregulation in the Δ*efhP* response to Ca^2+^. Interestingly, by 12 h Ca^2+^ exposure, transcription of most pyoverdine genes in Δ*efhP* returned nearly to the levels observed at no added Ca^2+^, but the pyochelin biosynthesis and uptake genes remain significantly down regulated. The Fe sequestering siderophores pyoverdine and pyochelin are known to serve critical roles in Fe acquisition from the host during *Pa* infection and contribute to virulence of the pathogen (*55–57*). The Has and Phu heme uptake systems are required for utilization of heme and playing a particularly significant role in *Pa* chronic infections of the CF lung (*58, 59*). Receptors PfeA and PirA have been established as essential for *Pa* uptake of the *Escherichia coli* siderophore enterobactin, further diversifying mechanisms of iron acquisition in this pathogen (*60*).

An opposite transcriptional profile was observed for the bacterioferritin gene *bfrB* and probable bacterioferritin PA4880, with these genes being downregulated both rapidly and adaptively in PAO1 in response to Ca^2^ but upregulated in Δ*efhP.* This stands consistent with the general Fe starvation transcriptional response (*61*) observed in response to Ca^2+^ in PAO1, as *bfrB* and PA4880 have previously been identified as part of Fe uptake master regulator Fur’s positive regulon, showing induction under growth at high Fe conditions (*61, 62*).

To sense and respond to environmental stimuli, PAO1 encodes 19 extracytoplasmic function sigma factors (ECFσ) (*63*). The ECFσ systems typically include transmembrane anti-σ sensors and cytoplasmic σ factors, which function to tightly regulate *Pa* stress responses (*64*). Interestingly, a majority of identified ECFσ in PAO1 (14 of 19) have been shown to respond to Fe starvation (*63*). We observed a major transcriptional induction of the Fe starvation ECFσs by Ca^2+^ in PAO1, which requires EfhP (Fig. 5A).

While transcription of the global regulator of iron uptake, *fur,* showed no significant Ca^2+^-responsiveness in PAO1, in Δ*efhP,* we observed 1.8-fold and 2.2-fold upregulation of *fur* following 60 min and 12 h exposure to Ca^2+^, respectively (Fig. 4A). These results were validated via RT-qPCR measurement of *fur* transcripts in PAO1 and Δ*efhP* (Fig. 4B). In agreement, according to RNA seq, the transcription of Fur-repressed *pvdS* and *pchR*, encoding important regulators of pyoverdine and pyochelin production, respectively (*65, 66*), was upregulated in PAO1 following 10 min, 60 min, and 12 h exposure to Ca^2+^ but downregulated in Δ*efhP*.

**Figure 4B.**
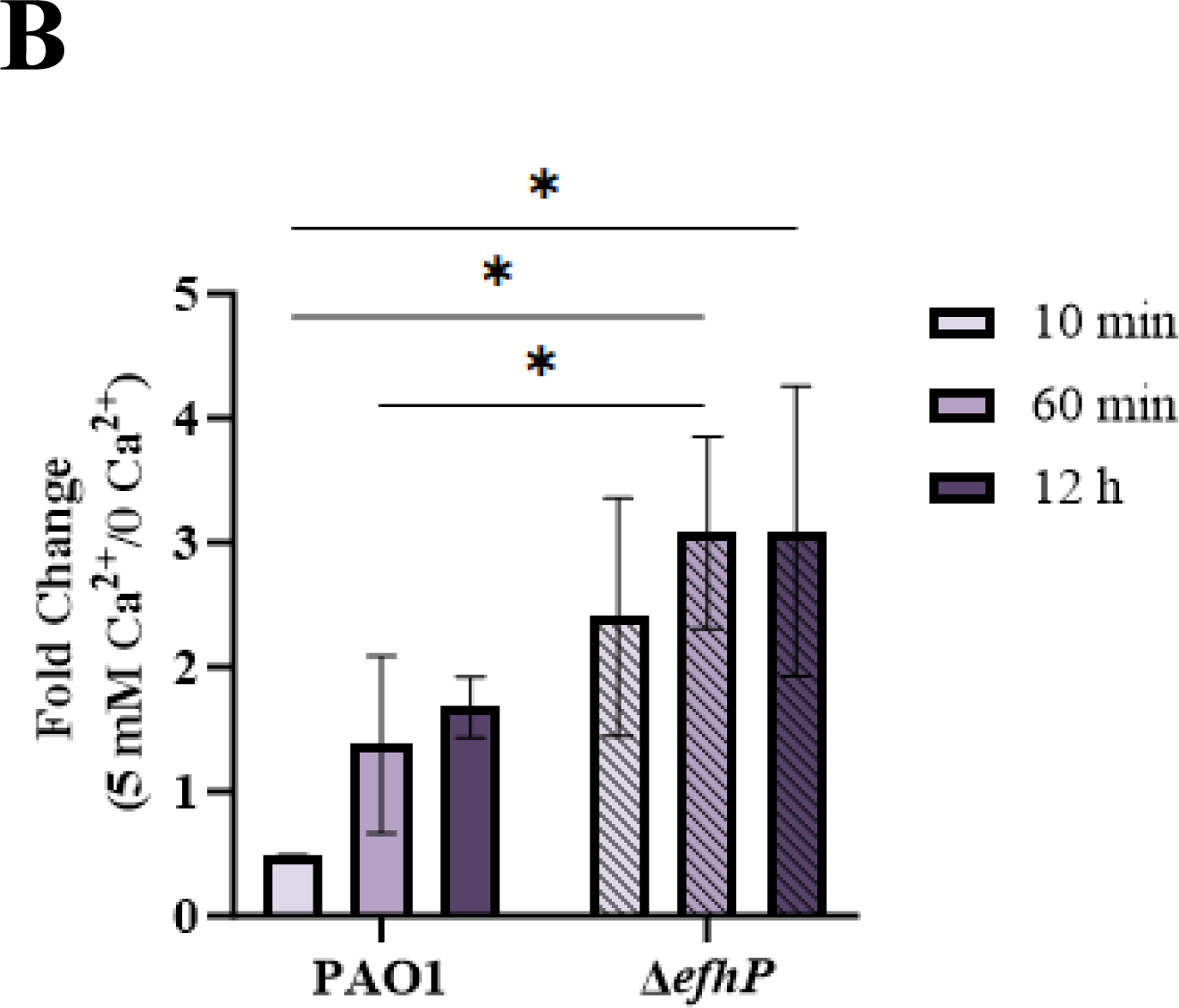
Impact of Ca^2+^ on transcription of *fur* in PAO1 and Δ*efhP*. qPCR was used to evaluate the changes in *fur* transcription in mid-log PAO1 and Δ*efhP* mutant cells grown in BMM at 10 min, 60 min, or 12 h 5 mM Ca^2+^ conditions vs. 0 mM Ca^2+^ condition. The relative mRNA levels were normalized to those of *rpoD*. Fold difference in relative transcription at 5 mM *vs* no Ca^2+^ was calculated using the 2^-ΔΔCt^ method. Statistical significance was determined using an unpaired one-tailed t-test, * indicates *p* < 0.05.

To validate EfhP regulation of *Pa* iron uptake, we have assayed for pyoverdine production in PAO1, Δ*efhP* and Δ*efhP::efhP* cultures. Supporting the transcriptional data, stationary phase (24 h) PAO1 showed a 7.8-fold increase in pyoverdine production when cultured at 5 mM Ca^2+^ compared to the 0 mM Ca^2+^ condition (Fig. 6A). This Ca^2+^ regulation of pyoverdine induction was lost in Δ*efhP* but recovered with *efhP* complementation. Since this response was observed in the cultures grown in the presence of 3.6 μM Fe in the medium, next, we aimed to characterize the impact of elevated Ca^2+^ on pyoverdine production in PAO1, Δ*efhP,* and Δ*efhP::efhP* cultures under iron starvation. Due to difficulties achieving consistent growth under prolonged iron starvation conditions, 3.6 μM added Fe was present in precultures, and main cultures were prepared in medium with no added Fe. Interestingly, when the cultures were starved for Fe, PAO1 and Δ*efhP* pyoverdine production showed to be insensitive to Ca^2+^ concentration (Fig. S1A), suggesting that the lack of essential Fe is higher in the hierarchy of signals than Ca^2+^. Nevertheless, pyoverdine levels normalized by OD_600_ in Δ*efhP* at both 0 mM and 5 mM Ca^2+^ concentrations were significantly lower than those observed in the wild type, indicating that EfhP is needed for a full scale of pyoverdine production under these conditions. Further, Δ*efhP* growth at no added Fe was stronger than that of PAO1 and Δ*efhP::efhP* at both 0 and 5 mM Ca^2+^ (Fig. S1B), possibly due to the reduced metabolic burden of pyoverdine production in this strain. At 3.6 μM Fe, all three strains showed stronger growth at 5 mM Ca^2+^ compared to 0 mM (Fig. S1B). While Ca^2+^-dependent Fe uptake may partially explain the improved growth performance in the presence of Ca^2+^ and Fe, the cause of continued growth advantage of Δ*efhP* over PAO1 and Δ*efhP::efhP* remains unknown.

**Figure 5.**
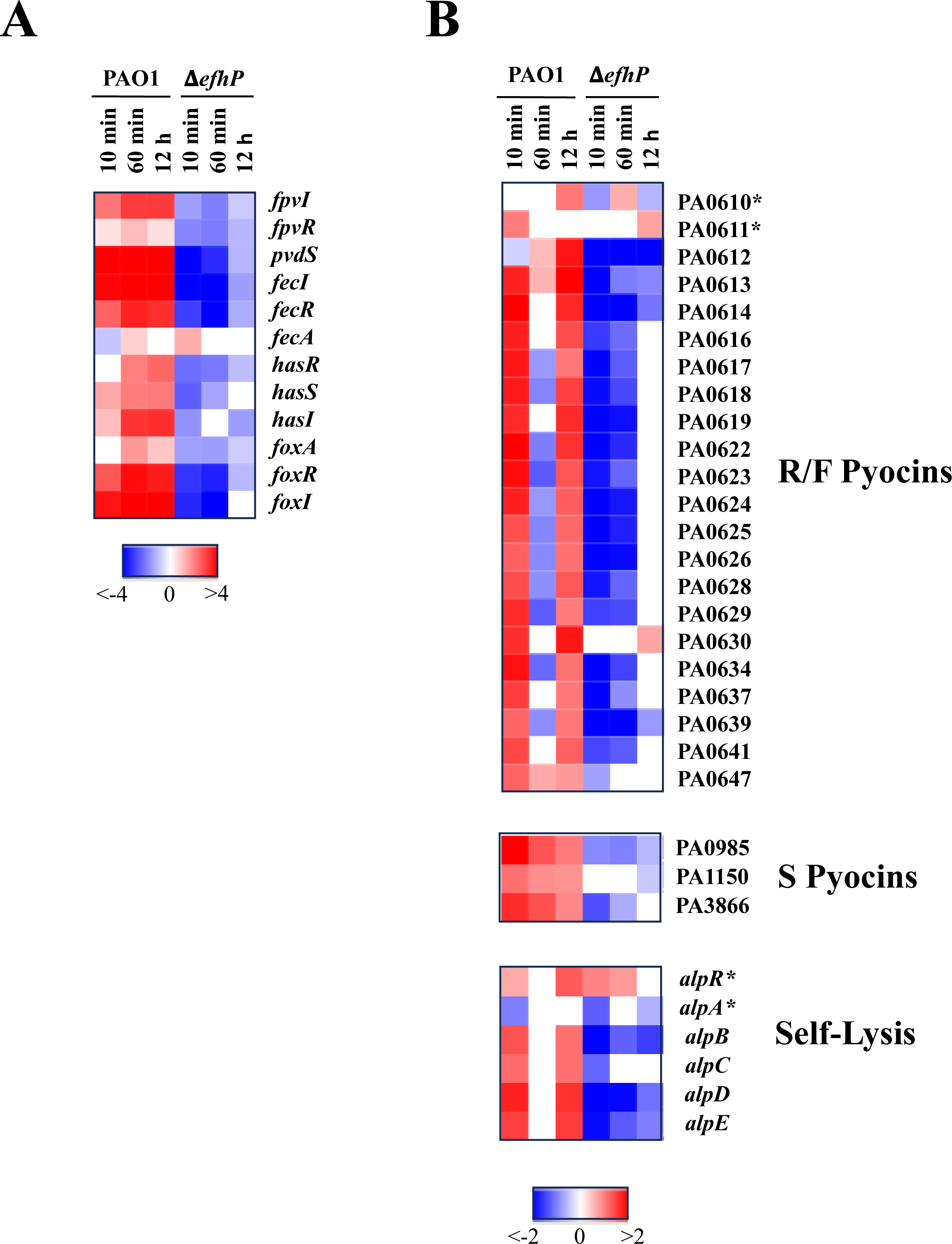
EfhP regulation of Fe starvation ECFs (A) and pyocin and self-lysis genes (B). The heat maps show log2FC values of expression comparisons of 10 min, 60 min, or 12 h 5 mM Ca^2+^ conditions vs. 0 mM Ca^2+^ condition. PAO1 and Δ*efhP* strains are compared to their respective control transcriptional profiles of cultures grown to mid-log phase at 0 mM Ca^2+^. ECF genes related to Fe starvation, as reviewed by Chevalier et al. 2019 are shown. Pyocin and self-lysis regulators are denoted with asterisks.

**Figure 6.**
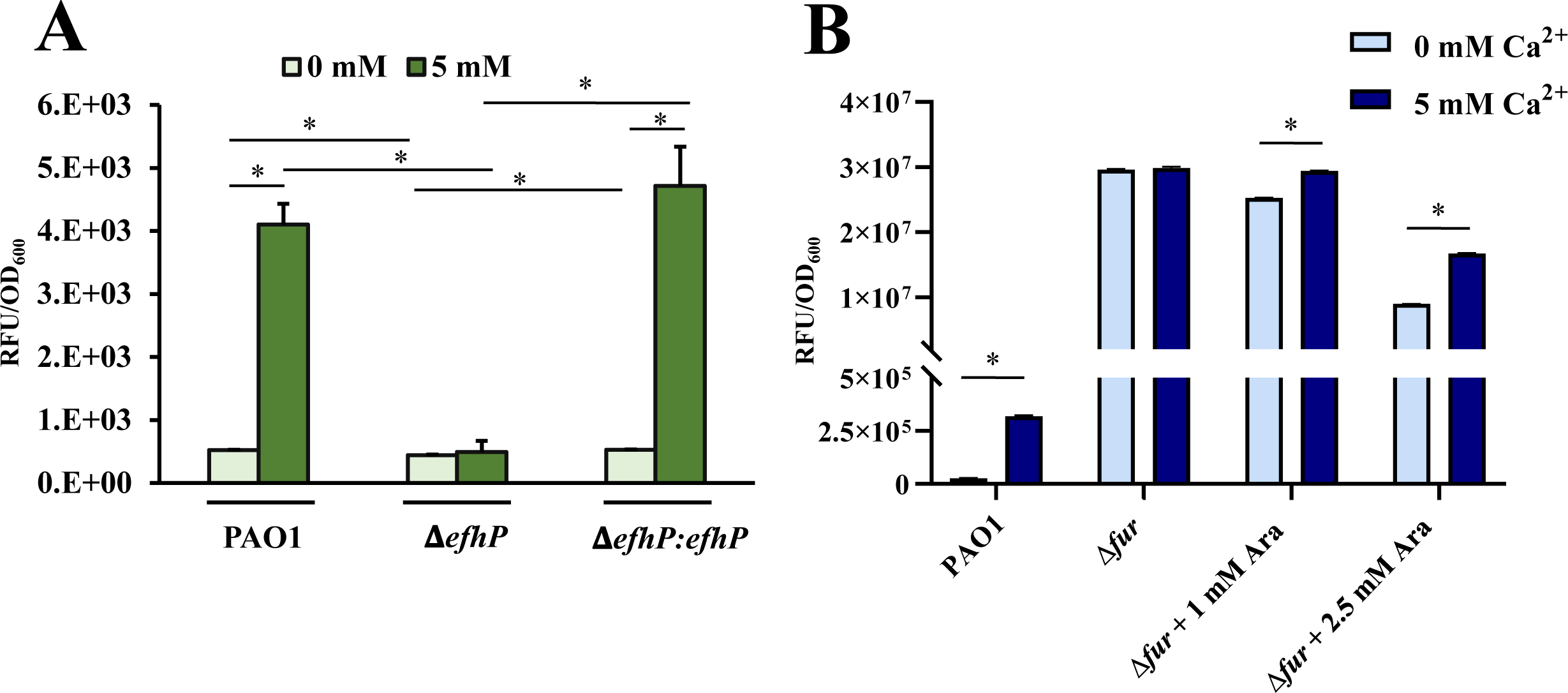
EfhP (A) and Fur (B) regulation of pyoverdine production at elevated Ca^2+^. Pyoverdine was quantified by reading fluorescence at 400 nm excitation/460 nm emission and normalized by OD_600_. (**A**) Stationary (24 h) phase pyoverdine production of PAO1, Δ*efhP*, and Δ*efhP::efhP* grown in BMM8 with or without 5 mM Ca^2+^ is shown as labeled. **(B)** Arabinose-dependent conditional *fur* deletion mutant (generously shared by Dr. Imperi) was cultured in the presence of 0 mM, 1 mM, or 2.5 mM arabinose to achieve increasing expression of Fur. Area under the curve was calculated to present total fluorescence produced during the time period from early log to after the transition to stationary phase. Statistical significance was determined by single factor ANOVA (* indicates *p* < 0.05).

To determine whether elevated Ca^2+^ leads to increased consumption of Fe causing an actual Fe starvation, we have quantified the remaining Fe in PAO1 and Δ*efhP* culture supernatants by inductively coupled plasma optical emission spectroscopy (ICP-OES). The presence of Ca^2+^ in growth medium during 10 m, 60 m, or 12 h exposure did not cause any significant reduction in Fe levels in either strain when compared to those observed at no added Ca^2+^ (Fig. S2A), supporting that Fe uptake is not increased at higher levels of Ca^2+^. It is noteworthy that the supernatants of Δ*efhP* at all Ca^2+^ conditions contained elevated Fe compared to those of PAO1, indicating that EfhP contributes to full Fe uptake in these conditions. We also investigated whether Ca^2+^ binds pyoverdine and thus limits its availability for binding Fe. Spectral scans of pyoverdine-rich filtrates incubated in the presence of 5 mM Ca^2+^ showed slightly less pyoverdine available to bind Fe compared to 0 mM Ca^2+^ condition (Fig. S2BC) and suggested that pyoverdine has a low affinity for Ca^2+^ as has been observed for numerous other metals (*67*). However, this limited effect does not seem to account for the significant changes in pyoverdine production seen at high Ca^2+^.

Finally, since Ca^2+^ regulates the entire Fe acquisition system in *Pa,* which is controlled by the negative regulator Fur (*68*) we hypothesized that Fur may be involved in this regulation. To test this hypothesis, we used *fur-*conditional mutant (*69*) that expresses Fur upon induction with arabinose. We showed that when Fur is absent (at no arabinose added), the production of pyoverdine increased about 1,000-fold *vs* that in the WT at no Ca^2+^ (Fig. 6B). This agrees with its negative regulation of *pvd* genes. In addition, we observed no Ca^2+^ induction in the mutant. However, when increased amounts of arabinose were added to the cultures (1 mM and 2.5 mM), the level of pyoverdine decreased in the presence of Ca^2+^ and thus, the induction of Ca^2+^ was restored (Fig. 6B), These results suggest that Ca^2+^ regulation of pyoverdine production involves Fur. The mechanisms of this involvement will be the focus of future studies.

### EfhP is required for Ca^2+^ regulation of pyocins and self-lysis

EfhP’s regulation of the *Pa* response to Ca^2+^ extends to pathways involved in pyocin production and self-lysis (Fig. 5B). The *Pa* genome encodes three different types of bacteriocins (known as pyocins), which have bactericidal activity against same or closely related *Pseudomonas* species (*70*). This includes the R- and F-type pyocins, whose genes are clustered within a chromosomal locus in PAO1, and the S-type pyocins, which have a more scattered distribution across the genome (*70*). In this study, upregulation in PAO1 transcription of R- and F-type pyocins was observed following 10 min and 12 h exposure to Ca^2+^, while S-type pyocins were upregulated at all three conditions. This Ca^2+^-dependent induction of most pyocin genes was nearly absent in Δ*efhP* at all time points. Additionally, most of the *alp* self-lysis genes were upregulated in the PAO1 response to Ca^2+^ at 10 min and 12 h but downregulated in Δ*efhP.* Interestingly, transcription of regulators *alpR* and *alpA* was unaffected in the Δ*efhP* response to Ca^2+^, but Ca^2+^-exposure repressed much of their regulon, *alpBCDE*.

To validate the observed Ca^2+^ regulation of pyocin biosynthesis, we have tested the impact of Ca^2+^-induced pyocin production on the survival of R-type pyocin-sensitive *Pa* strain 13S (*71*). For this, the 13S strain was grown to middle log in BMM8, and the cultures normalized to OD_600_ of 0.3 were treated with the supernatants obtained from the PAO1, Δ*efhP,* and Δ*efhP::efhP* grown in the presence or absence of 5 mM Ca^2+^. The calculated fold differences in survival of 13S upon treatments with Ca^2+^-induced and Ca^2+^-non-induced supernatants of the three strains are shown in Fig. 7 In agreement with the transcriptomics data, the presence of Ca^2+^ during PAO1 growth resulted in the supernatants that reduced 13S survival 2 times (fold change 0.5). This relationship was inversed in the case of Δ*efhP,* where we observed 1.6 fold increased survival, and reversed back to the WT level in Δ*efhP::efhP.* These data support the RNA-seq data suggested Ca^2+^ induction of R-type pyocin production in PAO1, which requires the presence of EfhP.

**Figure 7.**
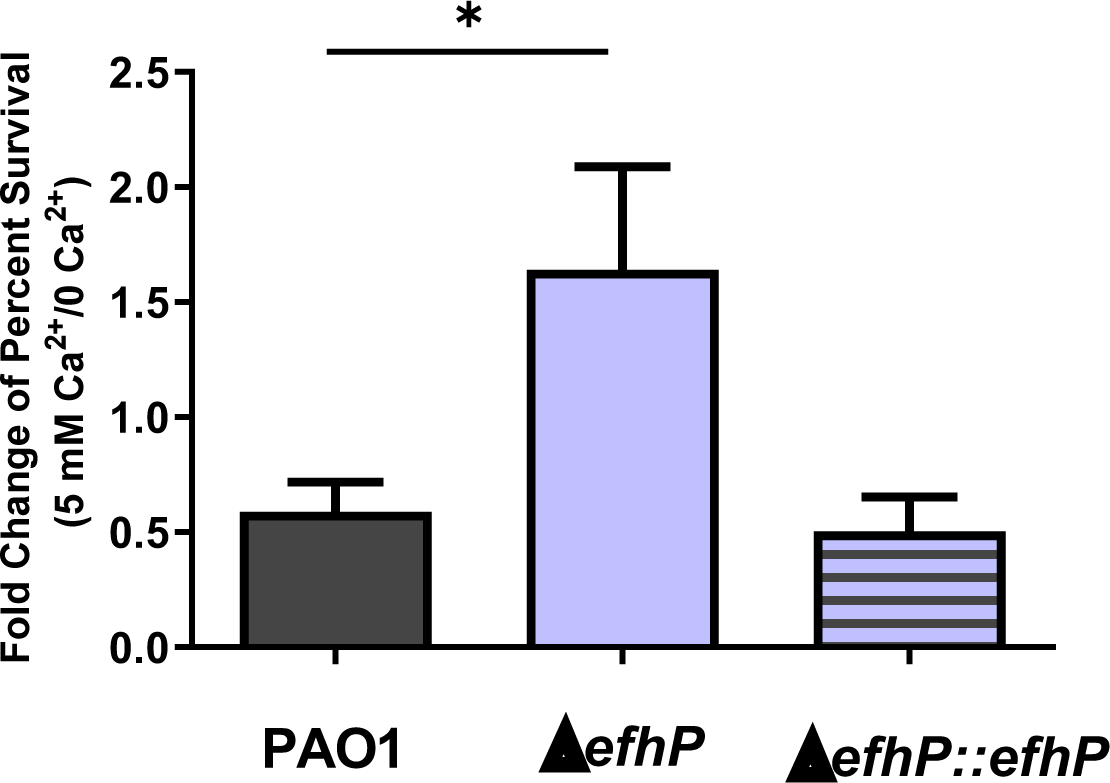
The impact of Ca^2+^-induced pyocin production on the survival of the *P. aeruginosa* R-type pyocin-sensitive strain 13S. Fold change of survival of 13S, upon addition of respective supernatants from 5 vs 0 mM Ca^2+^ cultures of wild-type PAO1, Δ*efhP*, and Δ*efhP::efhP*. 100 µL of OD_600_ 0.3 13S culture was treated with 100 µL of supernatants collected from PAO1, Δ*efhP*, and Δ*efhP::efhP* grown in 0 or 5 mM Ca^2+^ BMM8. The fold change in 13S survival as an effect of Ca^2+^ was calculated by dividing percent survival at 5 mM Ca^2+^ by that at no Ca^2+^. Treatments were performed in triplicate, and the experiments were repeated independently three times. An unpaired one-tailed t-test was used to determine statistical significance, * indicates *p* < 0.05.

### The expression of *efhP* is regulated by Ca^2+^, Fe, growth phase, and the two-component system CarSR

The RNA-seq analysis revealed that the expression of *efhP* was slightly increased after 12 h of PAO1 growth in the presence of Ca^2+^ but was not responsive to the ion after shorter exposures. Since this change was not statistically significant, we aimed to generate an additional insight, and tested the impact of 2 mM, 5 mM, and 10 mM Ca^2+^ on *efhP* expression during different phases of growth by using a promoter activity assay. It was observed that the presence of Ca^2+^ caused an increase in transcription of *efhP* during both the early log and stationary phases of growth but not in the mid-log phase (Fig. 8A).

**Figure 8.**
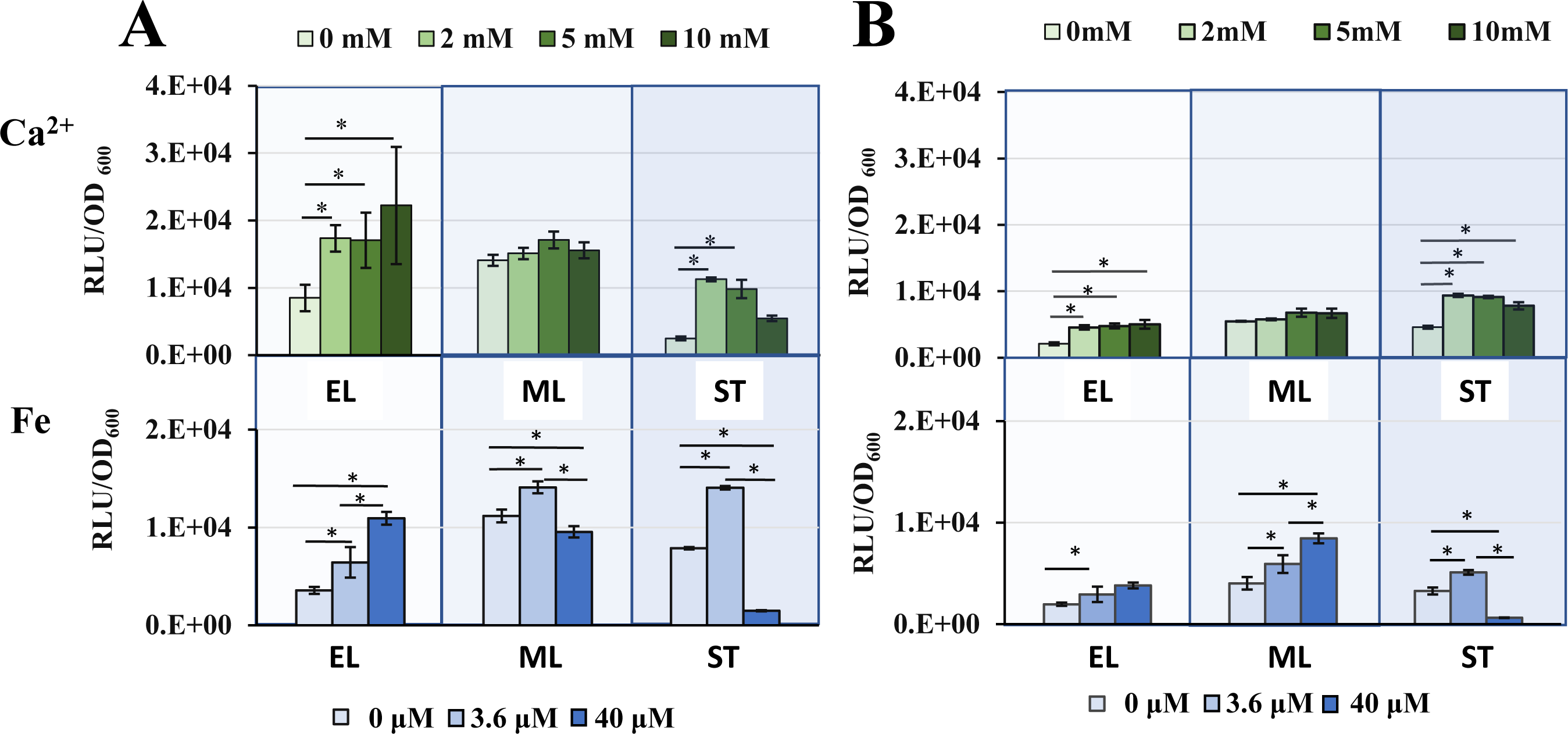
Impact of elevated levels of Ca^2+^ and Fe on *efhP* transcription. The wildtype PAO1 **(A)** or Δ*carR* **(B)** harboring the *efhP* promoter construct were grown at none or elevated levels of Ca^2+^ or Fe. Luminescence normalized by OD_600_ are shown for early log (4 h, EL), mid log (6-7 h, ML) and stationary (12 h, ST) for each strain and condition. Graphs represent the mean of at least three biological replicates. Error bars indicate the standard error. Statistical analysis was done by univariate ANOVA in SPSS (version 29.0.0.0, IBM Corp.2022) with a confidence interval of 95% (* indicates *p* < 0.05).

Considering the discovered role of Ca^2+^ in regulating Fe uptake, we also aimed to characterize the potential role of Fe in regulating the transcription of *efhP.* For this, we evaluated the promoter activity of *efhP* during growth of PAO1 at no added or provided with 3.6 μM or 40 μM Fe in the absence of Ca^2+^. Although the overall impact was smaller than that in response to Ca^2+^, the *efhP* promoter activity was also regulated by Fe (Fig 8A). However, in this case, we observed an increase in response to 3.6 μM at all the time points, and a decrease at 40 μM Fe in both mid-log and stationary phase cells.

Others have reported strong regulation of *efhP* transcription by the two-component system BfmRS, which was also shown to control *Pa* biofilm production (*72, 73*). We have chosen to investigate regulation of *efhP* by another *Pa* two-component system, CarSR (*48*). Our previous work showed that CarSR is regulated by Ca^2+^ and positively regulates the expression of several genes, including *carO* and *carP* in a Ca^2+^ dependent manner (*74*). An identical homolog of CarR in PA14, BqsR has been shown to be regulated by Fe (*75*). This indicates that CarR may mediate the regulatory impact of elevated levels of Ca^2+^ and Fe on transcription of *efhP_._* Hence, we measured the *efhP* promoter activity in the Δ*carR* mutant in response to elevated Ca^2+^ and Fe. Supporting the regulatory role of CarR, promoter activity of *efhP* in Δ*carR* background was significantly lower than that in PAO1 at most of the tested conditions (Fig 8B) with the strongest difference during early log phase.

### *efhP* expression is elevated in CF clinical isolates

It is important to understand whether increased *efhP* expression plays a role in *Pa* adaptation to the CF lung. According to the reported microarray data, the transcription of *efhP* is greatly increased in clinically relevant conditions. For example, *in vivo* microarray study utilizing sputum from the lungs of CF patients detected that *efhP* belongs to the top 1% of the most highly expressed *Pa* genes during infection (*76*). Additionally, *efhP* expression was shown to be greatly elevated in a CF clinical isolate exhibiting mucin sulfatase activity (*77*) as well as in later stages of infection with *Pa* (*78*).

To evaluate the levels of *efhP* expression in CF clinical isolates, we selected sputum *Pa* isolates from CF patients ranging in age from 7 to 55 years and tested for *efhP* expression via RT-qPCR. The different ages were selected for potential correlation between the level of *efhP* expression and the advancement of disease. Most CF isolates showed at least two-fold increased *efhP* expression in comparison to that in PAO1, with one of them (CF4) expressing *efhP* at levels 1,354 fold above those of PAO1 (Fig. 9A). While no clear trends in *efhP* expression emerged in relationship with the ages of CF patients, a larger sample of isolates may aid this effort in future studies.

**Figure 9.**
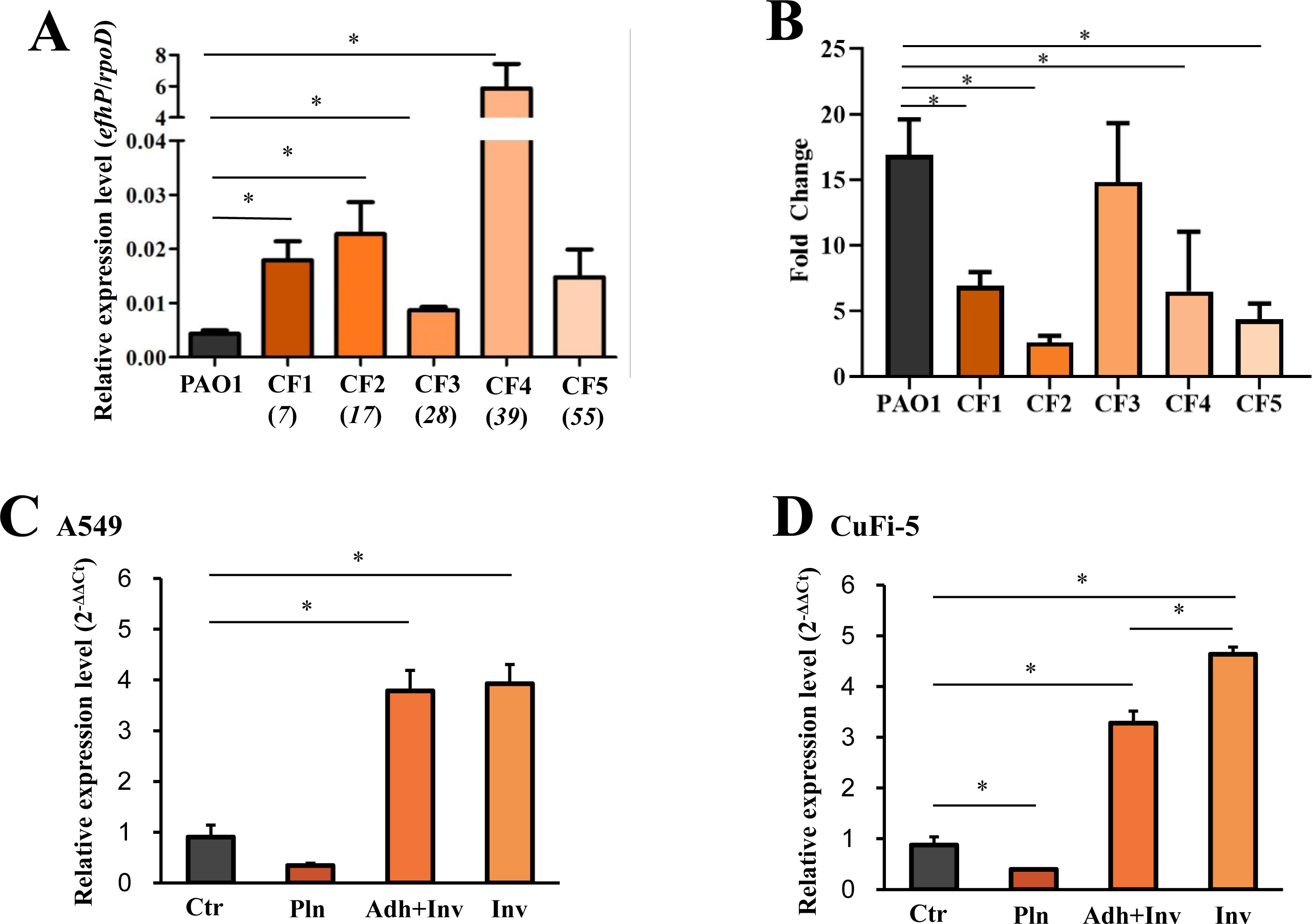
Analyses of *efhP* regulation (A) and pyoverdine production (B) in CF clinical isolates and *efhP* regulation during *Pa* interactions with A549 epithelial cells (C) and CuFi-5 epithelial cells (D). (A) Mid log *efhP* transcription by five CF clinical isolates grown in LB medium was compared to that of PAO1 by RT-qPCR. Ages of CF patients from which each isolate originated is indicated in italics. The relative mRNA levels were normalized using standard curve described in the methods. All significance between clinical isolates vs. PAO1 determined by single factor ANOVA (* indicates *p* < 0.05). **(B)** To measure pyoverdine production in CF clinical isolates, the strains were grown in BMM8 with or without 5 mM Ca^2+^. Pyoverdine was quantified by reading fluorescence at 400 nm excitation/460 nm emission and normalized by OD_600_. Area under the curve was calculated to present total fluorescence produced during the time period from early log to after the transition to stationary phase. Statistical significance was determined by single factor ANOVA (* indicates *p* < 0.05). **(C)(D)** To quantify *efhP* transcription in the presence of A549 **(C)** or CuFi-5 **(D)** epithelial cells, RT-qPCR was conducted using RNA obtained from growing PAO1 without epithelial cells (Ctr), planktonic (Pln), adhered+invaded (Adh+Inv), or invaded (Inv) PAO1. The relative mRNA levels were normalized to those of *rpoD* using the 2^-ΔΔCt^ method, and the fold difference was calculated versus Ctr. Median average and standard errors from three biological replicates are shown. For statistical analysis, both one-way ANOVA and Tukey-Kramer tests with a *P* of < 0.05 (*) were performed.

Based on the observed high level of *efhP* expression in the selected set of *Pa* CF clinical isolates, we hypothesized that the pyoverdine production in these isolates is increased in response to elevated Ca^2+^. To test this hypothesis, five selected isolates were grown in the presence or absence of 5 mM Ca^2+^ and their pyoverdine production was evaluated as described in Methods. As expected, we observed a significant Ca^2+^-dependent increase in pyoverdine levels in all five isolates (Fig. 9B). These data substantiate the regulatory role of Ca^2+^ for Fe uptake in clinical strains.

### The expression of *efhP* is regulated by interactions with host cells

Considering the reported and observed high levels of *efhP* expression in clinically relevant samples, we hypothesized that the expression of the gene is elevated during *Pa* interactions with host cells. To monitor the regulation of *efhP* expression during interactions of PAO1 with host cells, two types of human lung epithelial cell lines (adenocarcinoma alveolar basal epithelial cells A549 and CF bronchial CuFi-5 epithelial cells) were infected with PAO1. Following the infection, two experimental and two control populations of PAO1 were collected for RT-qPCR. The experimental populations included PAO1 adhered to epithelial cells and those internalized/invaded them. Since adhered PAO1 cells could not be separated from invaded cells, this group was considered as a mixture. As controls, we used planktonic PAO1 (free-swimming above the epithelial cells) or bacteria incubated alone. The quantities of the collected bacteria in each category are summarized in Table S1. The results revealed at least 3-fold increase in *efhP* expression upon interaction with host cells (Fig. 9CD). The increase in the *efhP* transcription during invasion is stronger in CuFi-5 cells when compared to A549 cells.

## DISCUSSION

Previous studies have identified Ca^2+^ regulation of numerous *Pa* virulence factors (*33, 34, 79*) contributing to increased mortality in *in vivo* infection models (*35, 49, 80*). The EF hand Ca^2+^ sensor, EfhP, was shown to control *Pa* Ca^2+^ responses including the induction of pyoverdine (*25*) and pyocyanin production (*49*) and elevated virulence in lettuce leaf and *G. mellonella* infection models (*49, 50*). In this study, we utilized RNA sequencing analysis to characterize the *Pa* rapid and adaptive transcriptional responses to Ca^2+^ which potentially drive the observed increases in virulence. Most notably, the results revealed a strong regulatory link between Ca^2+^ and Fe sequestering mechanisms. Numerous Fe uptake pathways in PAO1 were induced by Ca^2+^, including the biosynthesis and uptake of siderophores pyoverdine and pyochelin, and this regulation requires EfhP. Transcription of *efhP* itself is shown to be tuned by both Ca^2+^ and Fe, significantly elevated in *Pa* clinical isolates, and increased by interactions with host cells. These discoveries support the anticipated roles of EfhP in the *Pa* responses to the host-related factor Ca^2+^ benefiting its virulence and adaptation to the host.

Global RNA seq analysis indicated a capacity of PAO1 for kinetic transcriptional responses over the course from 10 min to 60 min to 12 h of Ca^2+^ exposure. While the Fe uptake genes account for a large share of Ca^2+^ responsive genes at all three time points, other functional categories made up more specific kinetic responses. These include sulfur metabolism and chemotaxis/motility genes in the rapid response to Ca^2+^, and secreted factors (pyocins, phenazines, toxins) and phosphate/phosphonate metabolism/transport genes in the adaptive response. The timing of these responses potentially reflects the functional significance of the corresponding processes for rapid and long-term adaptation strategies to the environments with elevated Ca^2+^. Interestingly, the 10 min condition shows the greatest number of differentially expressed genes, indicating that sensing Ca^2+^ triggers a massive transcriptional response and emphasizing the important regulatory roles of rapid Ca^2+^-sensing.

In addition to equipping *Pa* with Fe sequestering mechanisms, Ca^2+^ regulation of Fe homeostasis may also have implications for the multitude of Fe-controlled virulence factors. Production of the Fe^3+^-sequestering siderophores pyoverdine and pyochelin itself has been associated with increased *Pa* virulence (*55, 81, 82*). Further, production of biofilm and extracellular endoprotease PrpL (*33*) are regulated by both Fe (*83, 84*) and Ca^2+^, supporting the regulatory overlap of the signaling cascades (*33*). Co-regulation of virulence by Fe and Ca^2+^ may have especially significant impacts on pathogenicity of *Pa* in the CF lung where free Ca^2+^ (*51*) was shown to be elevated vs those seen in healthy individuals.

The RNA seq analysis showed that EfhP is required for 31% of the total PAO1 Ca^2+^ response, indicating the involvement of other Ca^2+^ signaling and regulatory systems, possibly including Ca^2+^-dependent two component system CarSR (*34*) and Ca^2+^ sensor LadS (*79*). Fe uptake pathways including pyoverdine, pyochelin, heme, and enterobactin as well as Fe storage fell under the regulatory control of EfhP at all the Ca^2+^ exposure intervals. The role of EfhP in regulating adaptive Fe uptake was supported by previous proteomic analysis conducted in PAO1 and mucoid strain FRD1 grown both planktonically and in biofilms (*25*). Deletion of *efhP* from these strains abolished or significantly reduced the Ca^2+^-dependent expression of several Fe uptake proteins, including pyoverdine biosynthesis proteins PvdNO, Fe(III)-pyochelin outer membrane receptor precursor FptA, ferric iron-binding periplasmic protein HitA, and a putative binding protein of iron ABC transporter PA5217 (*25*). In agreement, in the present analysis, all the encoding genes were transcriptionally regulated by EfhP at 5 mM Ca^2+^.

The observed Ca^2+^ induction of Fe uptake was not a result of Fe deficiency, as illustrated by ICP-OES-based total Fe quantification in PAO1 and Δ*efhP* strains cultured under the conditions used for RNA sequencing. The presence of elevated Ca^2+^ did not increase Fe uptake even after 12 h Ca^2+^ exposure. This supports the hypothesis that regulation of Fe uptake genes by Ca^2+^ is a result of Ca^2+^ signaling mechanisms rather than lack of Fe. Interestingly, Δ*efhP* showed significantly less Fe uptake at all Ca^2+^ conditions compared to PAO1. Taking into account the increased growth of Δ*efhP* at no Fe compared to PAO1, it appears that EfhP’s contributions to full Fe uptake became a growth hindrance when no Fe is present.

Another possible mechanism of Ca^2+^ interference with Fe-signaling cascades is via direct interaction with *Pa* siderophores. In addition to Fe^3+^, pyoverdine was shown to chelate 16 other metals at lower efficiencies, including Ga^3+^, Cu^2+^, and Ni^2+^ (*67*). Ga^3+^ disrupts *Pa* Fe homeostasis via competition with Fe^3+^ for binding to pyoverdine, from which it is unable to be reduced and released once in the periplasm (*67, 85, 86*). Cu^2+^, in addition to binding pyoverdine, was shown to stimulate pyoverdine production (*67, 87*), possibly via generation of oxidative stress. In this study, we show that 5 mM Ca^2+^ has a slight repressive effect on pyoverdine-Fe complex formation, likely due to a low pyoverdine affinity for Ca^2+^. While the ability of pyoverdine to bind Fe is partially hindered by elevated Ca^2+^, it appears that this impact is too subtle to cause Fe starvation.

In biological systems, Fe is present in one of its two distinct oxidation states, ferrous (Fe^2+^) or ferric (Fe^3+^) (*88*), with the ferric form being most abundant in aerobic conditions (*89*). *Pa* uses siderophores such as pyoverdine and pyochelin to bind Fe in its ferric state (*90*) which is reduced to its ferrous state and unloaded from pyoverdine in the periplasm. Once released from pyoverdine, Fe^2+^ enters the cytoplasm and binds ferric uptake regulator (Fur), which regulates genes involved in Fe homeostasis, uptake, and storage and thus controls the responses to the Fe availability (*88, 91*). It is currently unknown whether elevated intracellular Ca^2+^ concentrations, typically tightly regulated in *Pa* (*92*), have an effect on Fe^2+^ binding to Fur and other proteins. Deletion of *efhP* was shown to significantly disrupt intracellular Ca^2+^ homeostasis (*25*), providing a possible mechanism by which the Ca^2+^ sensor enacts regulatory control. While Fur is recognized as the master regulator of Fe uptake and is a conditionally essential protein in *Pa* (*69*), it does not act alone. Fur only indirectly regulates pyoverdine biosynthesis, acting as a repressor of σ factor *pvdS* under Fe-replete conditions (*93, 94*). When Fur binds Fe^2+^, it releases from the *pvdS* promoter region, allowing *pvdS* expression. PvdS then binds anti-σ FpvR, from which it is released upon PVD-Fe binding to receptor FpvA, resulting in transcription of the pyoverdine biosynthetic genes (*93, 95, 96*). EfhP’s predicted presence in the periplasm would allow it to interact with components of the Fe responsive ECFσ systems, such as FpvARI/PvdS, FecARI, HasRSI, and FoxARI which are transcriptionally regulated by Ca^2+^ in an EfhP-dependent manner as presented in this RNA seq analysis. One of the best studied periplasmic modulations of ECFσ activity is that by MucB, which binds MucA, preventing its proteolytic cleavage and subsequent release of σ AlgU (*97, 98*). Interaction between EfhP and an Fe starvation ECFσ at elevated Ca^2+^ could have strong effects on transcription of Fe uptake genes considering the high degree of positive feedback following activation of these systems (*99, 100*). We present a scheme of supporting evidence and current hypotheses for EfhP’s role in mediating the *Pa* Ca^2+^ response (Fig. 10).

**Figure 10.**
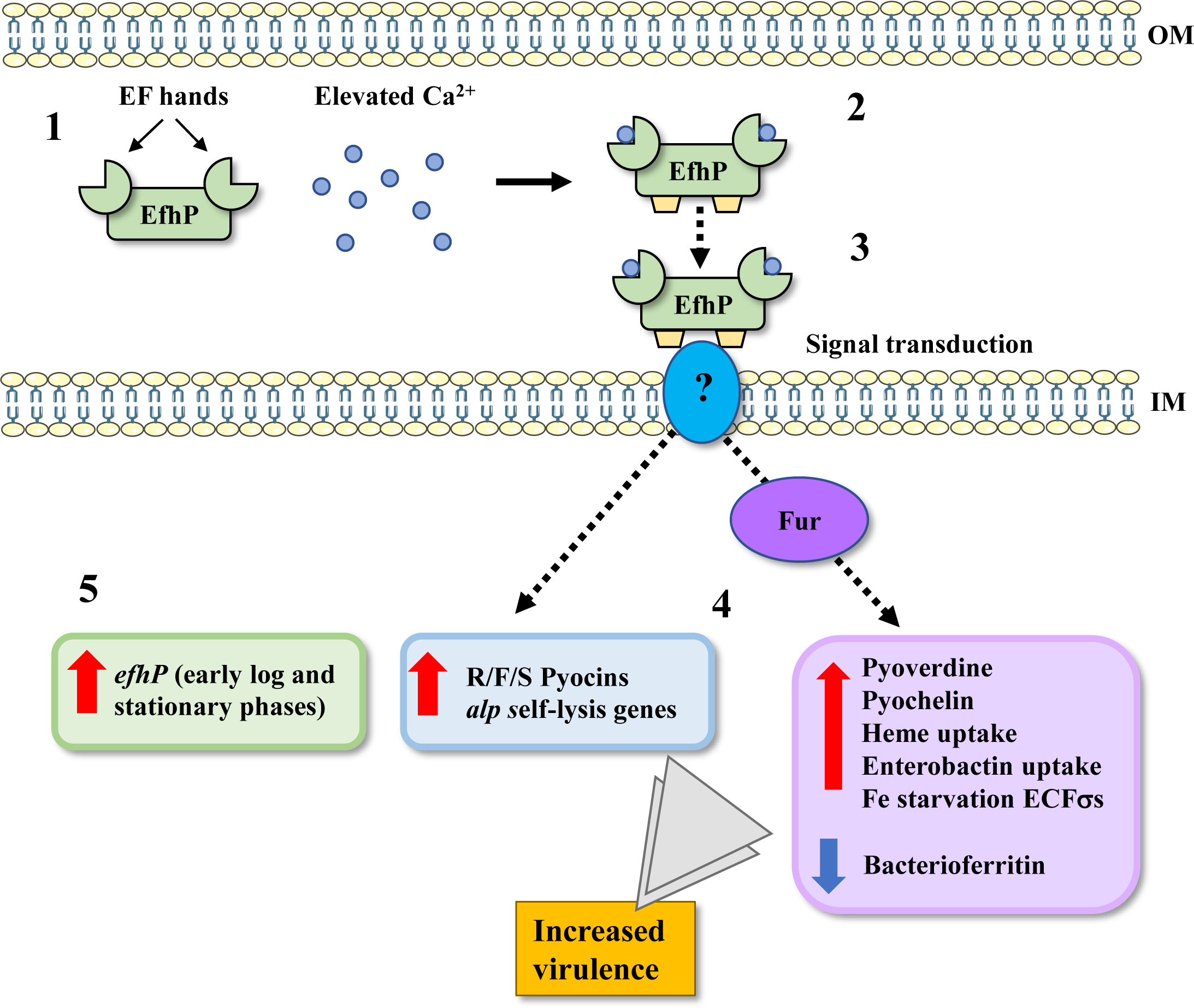
Role of EfhP in the *Pa* response to Ca^2+^. (**1**) Apo-EfhP is predicted to reside in the periplasm, where it can be exposed to elevated levels of Ca^2+^. (**2**) Upon binding Ca^2+^ with its EF hands, EfhP undergoes conformational changes, exposing hydrophobic regions of the protein (shown in yellow) (Kayastha et al. 2022). (**3**) It is hypothesized that these conformational changes allow EfhP to interact with still unknown protein partners and transduce the Ca^2+^ signals. (**4**) Rapid and adaptive transcriptional changes take place, as revealed by RNA sequencing analysis. Increases to the transcription of pyocin and self-lysis genes and the Fur regulon of Fe starvation genes are EfhP-dependent. Additionally, downregulation of select Fe storage genes (bacterioferritin) takes place at elevated Ca^2+^ in an EfhP-dependent manner. Together, this regulation of Fe starvation and pyocin/self-lysis by Ca^2+^ and EfhP is expected to increase *Pa* virulence in the host. EfhP’s role in Ca^2+^-induced *Pa* virulence has been previously established in lettuce leaf (Sarkisova et al. 2014) and *G. mellonella* (Kayastha et al. 2022) infection models. (**5**) Promoter activity data has revealed regulation of *efhP* transcription at elevated Ca^2+^ levels that is partially dependent on CarR. This suggests the presence of other *Pa* mechanisms for sensing of Ca^2+^, responsible for regulating the transcription of *efhP* and other EfhP-independent genes within the *Pa* Ca^2+^ response.

Direct sensing of Fe^3+^ or Fe^2+^ by EfhP is an additional possibility. The recent discovery of conserved cysteine residues within four repeats of ‘EGKCG’ in the EfhP sequence has led to the hypothesis that EfhP may serve dual Ca^2+^ and Fe-sensing functions. While these specific repeats have not yet been characterized, cysteine residues are known to serve Fe-sensing roles in other proteins (*101, 102*). One such example is the conserved Cys^675^ of *Pa* FeoB, which putatively binds Fe^2+^ and signals for the G-domain to bind GTP, opening the pore and allowing Fe^2+^ to pass (*103*). Dual Ca^2+^ and Fe-sensing by EfhP would help explain the high degree of crosstalk between the two signaling cascades observed in this study.

In this study we have identified transcriptional regulation of *efhP* itself by Ca^2+^ and Fe, which requires the two-component system CarSR (BqsSR). CF sputum contains about four times higher levels of Fe (*104*) and two times higher levels of Ca^2+^ (*51*) as compared to that in healthy individuals. Hence, it is possible that the elevated levels of Fe and Ca^2+^ in the host environment regulate the expression of *efhP* during the infection process, as was observed in the promoter activity data. While the presence of Fe promoted expression of *efhP* in early log phase, higher levels of Fe acted as a repressor of *efhP* expression in mid-log and stationary phases, suggesting additional growth phase-dependent regulation. Elevated Ca^2+^ had a positive regulatory effect on *efhP* expression in both early log and stationary phases. Increased transcription of *efhP* is thus identified as part of the *Pa* response to both Ca^2+^ and Fe, adding new dimension to its regulation of Fe uptake at high Ca^2+^ as revealed by the RNA seq analysis.

In this study, we also sought to further understanding of *efhP* expression during infection. Microarray expression data from the GEO database showed that the transcription of *efhP* is >150-folds higher in *Pa* isolates from CF sputa grown both *in vivo* and *in vitro* (GDS2869 and GDS2870), indicating the importance of this gene during infection and suggesting that its expression is responsive to the host environment. In this study, we showed that all five selected CF isolates express *efhP* at the levels higher than PAO1 *in vitro*. Additionally, we showed that expression of *efhP* is increased by *Pa* interaction with host epithelial cells. Upon invasion of host cells, *Pa* transitions from millimolar levels of Ca^2+^ in the extracellular space to nanomolar levels of free cytosolic Ca^2+^ (*30*). The transcription of *efhP* increases following invasion despite the lowered Ca^2+^, suggesting regulation by stronger, non-Ca^2+^ triggers and emphasizing EfhP’s role in *Pa* intracellular survival. Together with the previously demonstrated impact of EfhP on virulence in various infection models, it is expected that the Ca^2+^ sensor plays a significant role in *Pa* adaptation and virulence during infection of the human host.

## MATERIALS AND METHODS

### Bacterial strains, plasmids, and media

Table 1 lists the bacterial strains and plasmids used in this study. Biofilm minimal medium (BMM) (*39*) was used to grow *Pa* strains. BMM consists of 9.0 mM sodium glutamate, 50 mM glycerol, 0.02 mM MgSO_4_, 0.15 mM NaH_2_PO_4_, 0.34 mM K_2_HPO_4_, 145 mM NaCl, 200 μl trace metals, and 1 ml vitamin solution (per liter of medium). The trace-metals mixture was prepared with 5.0 g CuSO_4_.5H_2_O, 5.0 g ZnSO_4_.7H_2_O, 5.0 g FeSO_4_.7H_2_O, and 2.0 g MnCl_2_·4H_2_O per liter of 0.83 M concentrated hydrochloric acid. Vitamin solution was prepared by dissolving 0.5 g thiamine and 1 mg biotin (per liter of the final medium). Filter sterilized trace metals, MgSO_4_ and vitamin solution were added aseptically to autoclaved medium adjusted to a pH of 7. This prepared BMM has a final Fe concentration of 3.6 µM. To prepare ‘No Fe’ medium, FeSO_4_.7H_2_O was omitted from the trace metals. To encourage stronger growth in No Fe conditions where indicated, MgSO_4_ was added to a concentration of 0.08 mM (BMM8). When required, FeSO_4_.7H_2_O was added to the final concentration of 20 µM, 40 µM, 60 µM and 100 µM. When needed, 2, 5 or 10 mM CaCl_2_.2H_2_O (Sigma) was added. PAO1 was used to obtain genomic DNA for cloning *efhP.* For DNA manipulations, *E. coli* and *Pa* cultures were grown in Luria–Bertani (LB) broth (per liter: 10 g tryptone, 5 g yeast extract, 5 g NaCl) at 37 ^0^C with shaking at 200 rpm. Antibiotics used for *E. coli* were (per ml) 50 μg kanamycin (Kan) or 100 μg carbenicillin (Carb); for *Pa*, (per ml) 60 μg tetracycline (Tet). The plasmid pCTX-1-lux (*105*) was provided by Dr. Cabeen (Oklahoma State University). *E. coli* SM10 and *Pa* PAO1 were grown in LB broth shaking at 200rpm for 37°C or on LB agar plates with 1.5% agar at 37°C. For antibiotic selection plates, Tet was used at concentrations of 60 μg/ml (for *Pa*) and 10 μg/mL (for *E. coli*).

**Table 1.**
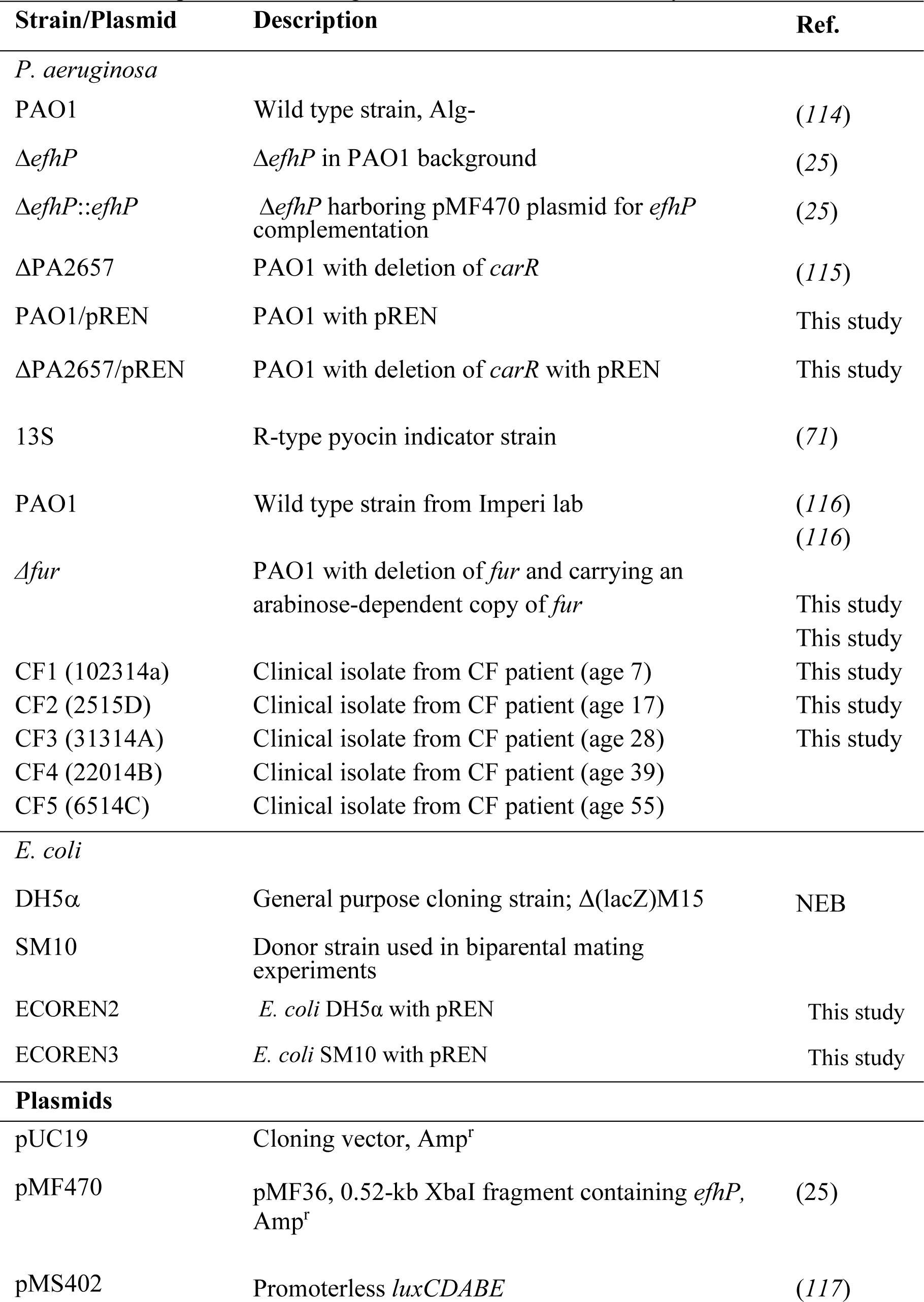
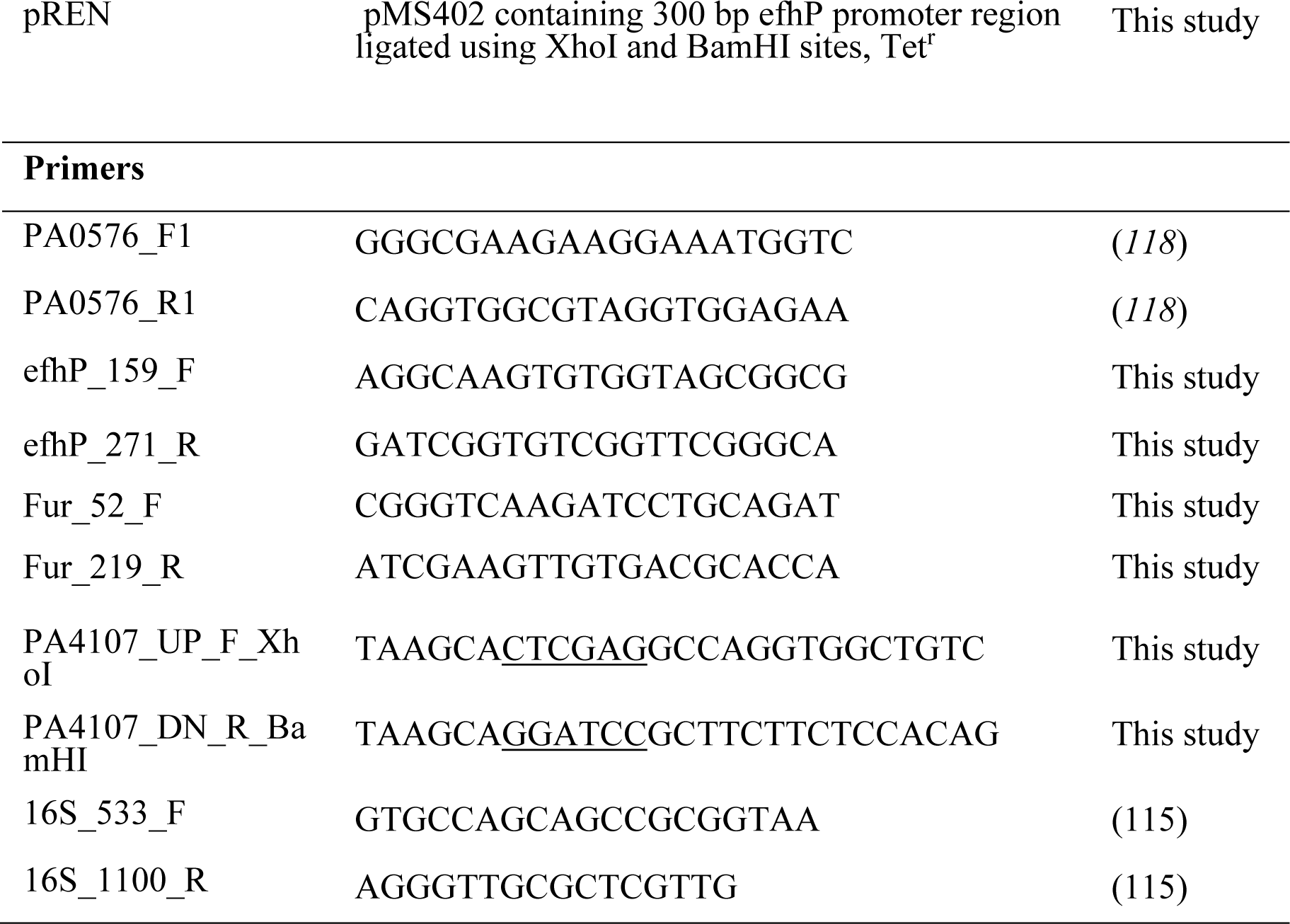
Strains, plasmids, and oligonucleotides used for this study.

### Standard DNA procedures

Plasmid DNA was isolated using the Zyppy plasmid miniprep kit (Zymo), and chromosomal DNA was obtained using the Wizard® Genomic DNA Purification Kit (Promega). DNA fragments were purified from agarose gels using the Zyppy gel extraction kit. DNA concentration was determined spectrophotometrically using NanoDrop 1000 Spectrophotometer (Thermo Scientific, Waltham, MA). DNA transformation into *E. coli* DH5α and *Pa* PAO1 was performed using heat shock or electroporation protocols.

### RNA sequencing

For RNA sequencing (RNA-seq) analysis, total RNA was isolated using RNeasy Mini Kit (QIAGEN) from *Pa* PAO1 cultures grown until mid-log phase in BMM either supplemented with 5 mM CaCl_2_, or without it to study adaptive response to Ca^2+^. To study rapid response to Ca^2+^, PAO1 cultures grown until mid-log with no CaCl_2_ were exposed to 5 mM CaCl_2_ for 10 or 60 min. DNase treatment was performed using turbo DNase (Ambion). The absence of genomic DNA was confirmed by conventional PCR using DreamTaq Green PCR Master Mix (Thermo Fisher Scientific) and primers 16s-533-F and 16s-1100-R targeting the 16S rRNA gene. RNA yield was measured using a NanoDrop spectrophotometer (Thermo Fisher Scientific), and the quality of the purified RNA was assessed with an RNA 6000 Nano Chip on a Bioanalyzer 2100 (Agilent) and by 1.2% agarose gel electrophoresis. Following the MIQE guidelines, only the RNA samples with an OD_260_/OD_280_ ratio of 1.8-2.0 and an RIN value of ≥ 7.0 were selected for further analysis. For sequencing, 250∼300 bp insert size, strand specific libraries were prepared and sequenced in Novogene Co., Ltd. Ribosomal RNA was removed using Ribo-Zero rRNA Depletion Kit (Illumina) and paired-end sequencing (2×150 bp) was done on the Nova Seq 6000 platform.

### Analysis of RNA seq reads

All raw RNA sequencing reads were uploaded to KBase (https://www.kbase.us/) for further processing. A minimum of two biological replicates were used for analysis of each strain and condition. Reads were paired and quality checked using FastQC-v0.11.5 and trimmed with Trimmomatic-v0.36. The trimmed reads were aligned to the genome with HISAT2-v2.1.0 and transcripts assembled using Cufflinks-v2.2.1. A differential expression matrix was created using DESeq2-v1.20.0 and the resulting raw gene count matrix was uploaded to iDEP.91 (http://bioinformatics.sdstate.edu/idep90/) for further analysis. Principle component analysis (PCA) and the desired condition comparisons for differential expression as generated on iDEP.91 were utilized for this study. The responses of PAO1 and Δ*efhP* to Ca^2+^ were compared in differential expression matrices. The differences in gene expression were presented as log2 of fold change (log2FC). Significance was determined by adjusted *p* values (*p*_adj) generated by DESeq2. The *p*_adj values below 0.05 were considered statistically significant. Heat maps of selected pathways were generated using Morpheus (https://software.broadinstitute.org/morpheus/). Sorting of the full PAO1 genome into functional categories was made possible using existing annotations from Gene Ontology (GO) (http://geneontology.org/), KEGG (https://www.genome.jp/kegg/annotation/), and the Pseudomonas Community Annotation Project (PseudoCAP) (https://www.pseudomonas.com/pseudocap). All RNA-seq reads utilized for this analysis are accessible in the NCBI Sequence Read Archive database (accession: PRJNA874094).

### Construction of promoter activity reporter

For studying the transcriptional regulation of *efhP*, a promoterless luxCDABE reporter was used. The pMS402 vector was generously provided by Dr. Kangmin Duan (Manitoba University, Canada). A 300 bp region upstream of *efhP* was amplified from the PAO1 genome by using Phusion polymerase and primers PA4107_UP_F_XhoI and PA4107_DN_R_BamHI (Table 1). The size of the PCR products was verified by agarose gel electrophoresis and purified by using the Zyppy extraction kit (Zymo). The amplified PCR products were digested with BamHI and XhoI and ligated with similarly digested pCTX-1-lux, upstream of the *luxCDABE* genes using the Quick Ligation kit (NEB). For transformation, the ligation mixture was added to heat shock competent DH5α *E. coli* cells and transformed using the protocol described above. The successful transformants were selected on LB agar plates containing 20 µg/μl Tet. The resultant plasmid construct confirmed by DNA sequencing, named pREN and was maintained in DH5α which was designated ECOREN2.

### Biparental mating for promoter activity constructs

The plasmid construct pREN was transformed into the mating strain *E. coli* SM10 by heat shock transformation and designated SM10REN2. It was then conjugated with PAO1 (WT) and the mutant Δ*carR* by biparental mating (*106*). The donor *E. coli* with the promoter construct and recipient strains were grown in 5 ml of LB overnight at 37°C. The recipient (1 ml) was incubated at 42°C for 2 h. The donor (50 µl) was spot dried on a LB plate in an aseptic zone for an hour. The heat-treated recipient (50 µl) was spot dried on top of the donor and incubated at 37°C for 24 h. The mating spot was scraped and suspended in 1 ml of phosphate buffered saline (PBS), out of which 10 µl was spread on a Pseudomonas Isolation Agar plate (PIA) supplemented with 60 µg/ml Tet and allowed to grow at 37°C, until colonies appeared. Resultant colonies were replica-plated on LB with 60 µg/mL Tet and grown at 37°C until colonies appeared. These colonies were tested for luminescence to verify transformation. Once verified, the strains were grown in LB with 60µg/ml Tet overnight and stored in 10% skimmed milk at -80°C. Empty pCTX-1-*lux* was transformed into PAO1 to be used as a control using biparental method as described above.

### Promoter activity assay

*Pa* strains carrying pREN were grown in 5 ml of BMM while shaking at 200 rpm for 12 h. OD_600_ of these cultures was measured and normalized to 0.3 by using sterile BMM. Then it was used to seed a main culture (200 µl final volume of BMM with desired concentration of Ca^2+^ and/or Fe) at a ratio of 1:100 in 96 well white clear bottom plates (VWR). The plates were covered with lids treated with a mixture of 95% ethanol and 1% Triton solution to prevent condensation. Then the plates were incubated in the Biotek plate reader at 37^0^C shaking at 200 rpm for 13 h programmed to measure the OD_600_ and luminescence in Relative Luminescence Units (RLU) every 1 h. The resultant RLU values were normalized by OD_600_. Values from non-inoculated cultures were used as blanks. Each experiment was conducted with at least three biological replicates and each experiment was conducted at least twice. Statistical analysis was done by univariate ANOVA in SPSS (version 29.0.0.0, IBM Corp.2022).

### Quantification of pyoverdine production

Intrinsic fluorescence of pyoverdine at 400 nm excitation/ 460 nm emission (*107*) was used to quantify pyoverdine production in PAO1, Δ*efhP*, and Δ*efhP::efhP* cultures. Twenty-four h colonies grown on LB plates containing appropriate antibiotics were used to inoculate precultures in 3 ml BMM8 with no Ca^2+^ and no Fe to deplete intracellular Fe. Following 12 h of growth, the cultures were normalized to an OD_600_ of 0.3, then diluted 1000x in BMM8 with or without 5 mM Ca^2+^, and 200 μl of the resulting cultures were transferred into black clear-bottom 96 well plates and incubated in BioTek SynergyMx at 37^0^C with shaking at 200 rpm. Fluorescence (400/460 excitation/emission, 9 nm bandwidth, gain= 50) and OD_600_ were measured hourly. Relative fluorescence units (RFU) were normalized by OD_600._

For quantification of pyoverdine production in cultures containing no added Fe, PAO1, Δ*efhP*, and Δ*efhP::efhP* precultures were grown in BMM8 (3.6 μM Fe) to ensure stronger growth. Twelve h precultures were normalized to an OD_600_ of 0.3 and diluted 1000x in BMM8 No Fe with or without 5 mM Ca^2+^. Hourly fluorescence and OD_600_ reads were taken in the BioTek SynergyMx as described above. For all pyoverdine assays, at least two independent experiments were conducted, each containing three biological replicates. For the cultures of clinical isolates and mutants with different growth patterns, pyoverdine production was calculated as area under the curve to present total fluorescence produced during the time period from early log to after the transition to stationary phase. Statistical significance was determined by single factor ANOVA (Microsoft Excel version 16.54) with cutoff of *p*< 0.05.

### Total Fe quantification by inductively coupled plasma optical emission spectroscopy (ICP-OES)

Samples for ICP-OES were obtained from PAO1 and Δ*efhP* cultures grown at 0 mM and 5 mM Ca^2+^ to mid-log phase (OD_600_ 0.2±.03) in 100 ml BMM. Following collection of 15 ml samples from the 0 mM cultures, 5 mM CaCl_2_ was added to the same flasks. Samples were subsequently taken again at 10 min and 60 min Ca^2+^ exposure. The 12 h Ca^2+^ condition utilized cultures grown in the presence of 5 mM Ca^2+^ to mid-log phase. All samples were spun at 6,010 g for 10 min at 4 ^0^C, then supernatants passed through 0.2 μm filter. Filtered supernatants were stored at -20 ^0^C until ICP-OES analysis. ICP-OES was performed using the iCAP PRO XPS ICP-OES Duo Spectrometer coupled with Qtegra ISDS software. A wavelength of 259.940 nm (Aqueous-Radial-iFR) was selected to detect Fe specifically, limiting interference, in samples. Measurements provided in parts per million were converted to μM for analysis. Three biological replicates were averaged, and standard error was calculated. Significance was determined by single factor ANOVA (Microsoft Excel version 16.54) with cutoff of *p*< 0.05.

### Pyoverdine spectral measurements

PAO1 was grown 16 h in 5 mL BMM8 (0 μM Fe, 0 mM Ca^2+^). Cultures were spun at 9,319 g for 10 min and filter sterilized to obtain pyoverdine-rich filtrate. Absorbance spectral scans of 1 mL filtrate samples were conducted in cuvettes in a BioTek SynergyMx. CaCl_2_.2H_2_O dissolved in H_2_O was added to a concentration of 5 mM in appropriate samples. Equal volumes of FeCl_3_ .6H_2_O dissolved in deionized H_2_O was added to filtrates at concentrations of 4 μM, 20 μM, 60 μM, and 1 mM FeCl_3_ following addition of either CaCl_2_ or H_2_O control. Deionized H_2_O without FeCl_3_ was added to 0 μM Fe samples. All samples were incubated 30 min at room temp before absorbance (Abs) reads. Raw Abs values were subtracted by BMM8 blank. Percentage bound pyoverdine was calculated using the pH insensitive specific pyoverdine-Fe Abs460 shoulder (*108*). Abs460 values for each sample were divided by Abs460 of Fe saturated pyoverdine (1 mM FeCl_3_, 0 mM Ca^2+^). Three technical replicates were conducted. Significance was determined by single factor ANOVA (Microsoft Excel version 16.54) with a cutoff of p< .05.

### *efhP* transcription during interactions with epithelial cells

Two types of human lung epithelial cell lines were used: adenocarcinoma alveolar basal epithelial cells A549 (ATCC® CCL-185) (*109*) and CF bronchial CuFi-5 (ATCC® CRL-4016) epithelial cells which are homozygous for the ΔF508 mutation of the CFTR protein in CF (*110*). A549 cell line in RPMI medium (ThermoFisher, MA) or CuFi-5 cell line in Airway Epithelial Cell Growth Medium (AECGM, PromoCell, Heidelberg, Germany) were seeded into six-well tissue culture plates and incubated at 37°C in 5% CO_2_ until reaching 100% confluence. PAO1 was grown in BMM until mid-log phase (12 h; OD_600_ =0.18±0.03). The bacterial cells were harvested and resuspended in the appropriate cell culture medium and inoculated to the apical cell surface at a multiplicity of infection (MOI) of 30. Cells were incubated for 2 h, extracellular bacteria were removed, media containing 200 μg/ml gentamicin (Gm) was added, and cells were incubated for an additional 4 h. For sampling of the invaded PAO1, cells were washed with PBS and lysed in 1 ml of ultrapure water. For sampling of the adhered bacteria, cells were incubated in the absence of Gm for 6 h, washed with PBS and harvested. This bacterial population also contained invaded bacteria. To count the bacterial cell number, the samples were 10x diluted and spotted onto LB agar plates, and CFU were calculated. For quantitative RT-PCR (RT-qPCR), total RNA was extracted as described above, and cDNA was synthesized using a Transcriptor first-strand cDNA synthesis kit (Roche) and RT-qPCR was performed using SYBR green master mix (Roche, Indianapolis, IN) and a Light Cycler 480 Instrument II (Roche) as described previously (*111*). The sequences of PCR primers used for RT-qPCR are provided in Table 1. The *rpoD* gene was used as an internal control. The relative fold changes in gene expression levels were calculated using the 2^-ΔΔCt^ method (*112, 113*).

### *efhP* transcription in CF clinical isolates

The clinical isolates from CF patients (age 7 to 55 years) were isolated from sputum samples collected from patients at the Children’s Hospital CF clinic in Oklahoma City, Oklahoma. Sputum samples were streaked on PIA and incubated at 37°C overnight until individual colonies appeared. Isolates were grown in liquid LB medium for 16 h and frozen stocks made in 10% skim milk before storage at -80 °C. To quantify *efhP* transcription via RT-qPCR, the strains were grown in LB medium until mid-log phase (OD_600_ of 0.30 ± 0.05). Total RNA was isolated, and cDNA was synthesized as described in (35). Absence of genomic DNA (gDNA) in the RNA samples was verified by PCR amplification of 16S *rRNA* genes prior to cDNA synthesis by using DreamTaq green PCR master mix, alongside a gDNA positive control. If gDNA was present, samples were run through a second DNase treatment.

Quantitative RT-PCR (RT-qPCR) was performed using SYBR green master mix (Roche, Indianapolis, IN) and a Light Cycler 480 Instrument II (Roche). Five serial dilutions of gDNA (10^1^-10^-3^ ng) were run as templates to plot a standard curve for *efhP* and *rpoD* primers. In a base-10 semi-logarithmic graph, the threshold cycle (Cp) values were plotted against the dilution factor, and an exponential trendline was fitted to the graph. Only a correlation coefficient (R^2^) greater than 0.99 was accepted for data analysis. The trendline equation was then used to determine relative expression levels of amplified genes. Expression of *rpoD* gene was used as a reference, and the relative *efhP* expression was calculated as the ratio *efhP/rpoD.* All primers used are listed in Table 1. Statistical significance was determined by single factor ANOVA (Microsoft Excel version 16.54) with cutoff p< 0.05.

### Validation of *fur* transcriptional regulation

Wild-type PAO1 was grown alongside Δ*efhP* in BMM with or without 5 mM CaCl_2_ to mid-log phase. Cells were harvested after approximately 12 h at an OD_600_ of 0.2±0.05. RNA was then isolated as described above, and cDNA was synthesized using a Transcriptor first-strand cDNA synthesis kit (Roche). Primers for *fur* (Table 1) were tested by qPCR to ensure amplification efficiency in relation to internal control *rpoD*. RT-qPCR was then performed using SYBR green master mix (Roche, Indianapolis, IN) and a Light Cycler 480 Instrument II (Roche) as described previously (*111*). The relative fold change in *fur* expression was calculated using the 2^-ΔΔCt^ method (*112, 113*). Statistical significance was determined by single factor ANOVA (Microsoft Excel version 16.54) with cutoff p< 0.05.

### Pyocin Assay

Wild-type PAO1, Δ*efhP*, and Δ*efhP::efhP* were grown in BMM8 with or without 5 mM CaCl_2_ to mid-exponential phase. Cultures were normalized to OD_600_ 0.3, pelleted, and filter sterilized. In parallel, overnight cultures of the R-type pyocin indicator strain, 13S, were normalized to OD_600_ 0.3. The supernatants of PAO1, Δ*efhP*, or Δ*efhP::efhP* (100 µL) were added to 100 µL of the normalized pyocin-indicator strain in 1.5 mL centrifuge tubes. The tubes were incubated for 30 min at 37℃. CFUs were determined following appropriate dilution and plating on LB plates. CFUs from before and after treatment were used to calculate percent survival. To determine the difference in survival as a Ca^2+^ effect, the fold change of percent survival was calculated by dividing percent survival at 5 mM Ca^2+^ by that at 0 mM Ca^2+^ condition. Treatments were performed in triplicates, the percent survivals were averaged and used to calculate fold difference between different Ca^2+^ levels. Each experiment was repeated independently three times.

### Data Availability

All raw RNA sequencing reads were uploaded to KBase (https://www.kbase.us/). All RNA-seq reads utilized for this analysis are accessible in the NCBI Sequence Read Archive database (accession: PRJNA874094)

---

**Figure S1. Growth and pyoverdine production of *Pa* at 0 µM added Fe. (A)** Stationary phase (24 h) pyoverdine production of no Fe BMM8 cultures of strains PAO1, Δ*efhP*, and Δ*efhP:efhP* as indicated was quantified by fluorescence at 400 excitation/460 emission and normalized by OD_600_. Statistical significance determined by single factor ANOVA (Microsoft Excel v16.54) with a *p* of < 0.05 (*). Strains PAO1 (black), Δ*efhP* (white), and Δ*efhP:efhP* (grey) were cultured in no Fe BMM8 in the absence (**B**) or presence (**C**) of 5 mM added Ca^2+^. Data represents hourly averaged OD_600_ values of three independent biological replicates.

**Figure S2. Total iron quantification of mid-log cultures (A) and spectral measurements of PVD-Fe complex formation (B-C). (A)** ICP-OES was employed to quantify total iron remaining in pelleted cultures of PAO1 and Δ*efhP* grown under same conditions as used for RNA sequencing. Data was converted from parts per million to µM for analysis. Significance determined by single factor ANOVA (p < 0.05 = *): a, significant against BMM control medium; b, significant between PAO1 and Δ*efhP* at 0 mM Ca^2+^ condition; c, significant between PAO1 and Δ*efhP* at 10 min Ca^2+^ condition; d, significant between PAO1 and Δ*efhP* at 12 hr Ca^2+^ condition. **(B)** Absorbance spectral scan of pyoverdine-rich filtrate in No-Fe BMM8 containing 0 µM (red), 4 µM (yellow), 20 µM (green), 60 µM (blue), or 1 mM (purple) added FeCl_3_. Scans were conducted in the presence (dashed) or absence (solid) of 5 mM CaCl_2_. **(C)** Percentage of bound pyoverdine following addition of each indicated FeCl_3_ conc. in the presence (dashed) or absence (solid) of 5 mM CaCl_2_. Abs_460_ of each sample was divided by the average Abs_460_ of the saturated 1 mM FeCl_3_ 0 mM CaCl_2_ samples to calculate percentages bound. Significance determined by single factor ANOVA (* indicates *p* < 0.05).

**Table S1.**
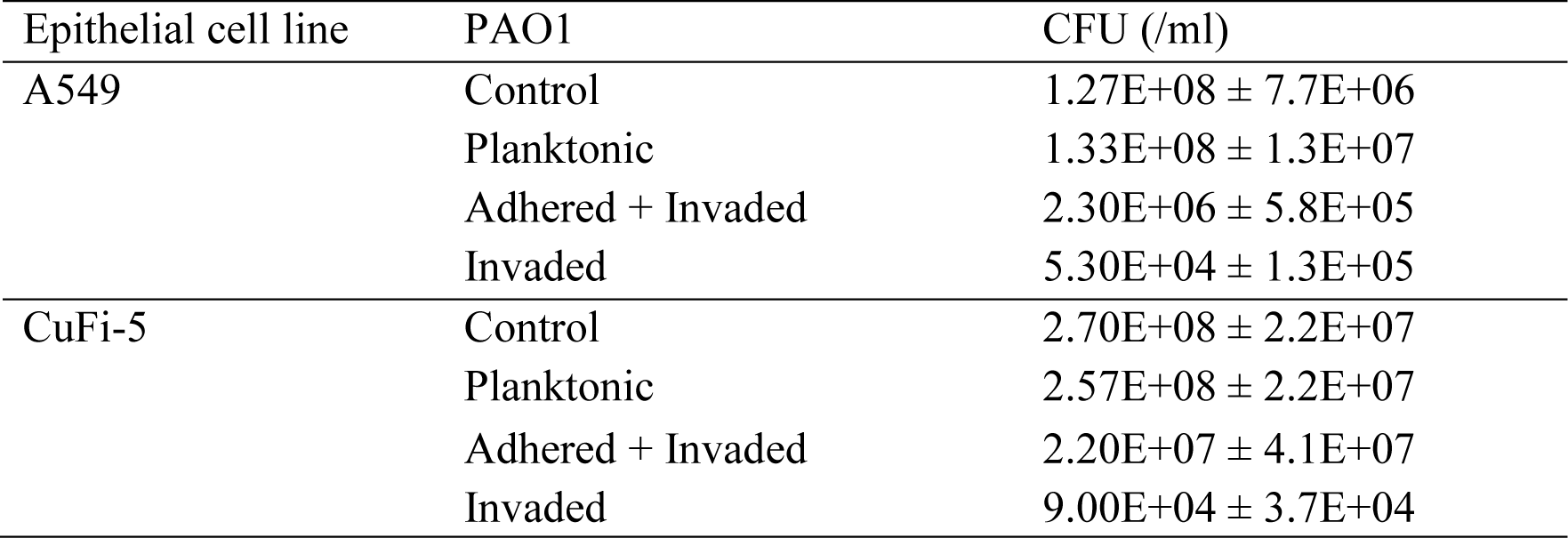
CFU results for PAO1 populations recovered after infecting A549 or CuFi-5 epithelial cells. Control condition reflects PAO1 grown in cell culture media described in Methods and in the absence of epithelial cells.

| Epithelial cell line | PAO1 | CFU (/ml) |
| --- | --- | --- |
| A549 | Control | 1.27E+08 ± 7.7E+06 |
|  | Planktonic | 1.33E+08 ± 1.3E+07 |
|  | Adhered + Invaded | 2.30E+06 ± 5.8E+05 |
|  | Invaded | 5.30E+04 ± 1.3E+05 |
| CuFi-5 | Control | 2.70E+08 ± 2.2E+07 |
|  | Planktonic | 2.57E+08 ± 2.2E+07 |
|  | Adhered + Invaded | 2.20E+07 ± 4.1E+07 |
|  | Invaded | 9.00E+04 ± 3.7E+04 |

## ACKNOWLEDGEMENTS

**MP** and **EL** were supported by NIH grant 1R15GM124670-0. MP was also supported by grant P20GM103648. We thank Dr. Chelsea Murphy at the Oklahoma State University formatic Center for her assistance with RNA seq plots. We are grateful to Dr. Francesco ri from Sapienza University of Rome for sharing the Fur conditional mutant.

